# Copper-containing ROS-scavenging nanozyme paradoxically drives α-synucleinopathy by amplifying nitrosative stress

**DOI:** 10.1101/2025.07.01.661317

**Authors:** Shuya Li, Yuanyang Mao, Ning Wang, Jiawei Zhang, Xinpeng Zhi, Yuyang Chen, Jingjing Si, Qi Yang, Ramhari Kumbhar, Deok Joeng, Ji-Yang Song, Xiuli Yang, Bong Gu Kang, Anika Khandekar, Janis Park, Yimeng Gao, Sean Yu, Rong Chen, Shu Zhang, Jingyan Han, Valina L. Dawson, Pengfei Liu, Shizhong Han, Ted M. Dawson, Weiwei He, Xiaobo Mao

## Abstract

Reactive oxygen and nitrogen species (RONS) are implicated in neurodegeneration, but their specific pathogenic roles remain unclear. Here, we developed a pair of iridium-based nanozymes with opposing functionalities to dissect these pathways. We show that a copper-tuned iridium nanozyme (Ir□Cu), despite being a superior ROS scavenger, paradoxically and dramatically exacerbated α-synuclein (αSyn) pathology *in vivo*. This pathology was causally linked to its ability to amplify RNS, as pharmacological inhibition of nitric oxide synthase (NOS) with L-NAME completely abrogated the pathology and reversed a human Parkinson’s disease (PD)-like transcriptomic signature. In contrast, a copper-free, broad-spectrum RONS-scavenging iridium (Ir) nanozyme demonstrated substantial therapeutic efficacy across diverse brain-first, body-first, and Alzheimer’s disease with Lewy body co-pathology models. Our findings uncover the importance of the RNS pathway in driving α-synucleinopathies and establish a critical design principle for nanomedicine, mandating caution in the use of redox-active copper for neuroprotective applications.

---

The aggregation of α-synuclein (αSyn) is the defining pathological hallmark of α-synucleinopathies, such as Parkinson’s disease (PD) and Lewy body dementia (LBD), and is also a significant feature in certain forms of Alzheimer’s disease (AD)^1–3^. A critical pathogenic feature of these devastating disorders is the prion-like propagation of misfolded αSyn aggregates^3–5^, often modeled using preformed fibrils (PFF)^6–13^, which drives the spread of pathology within the nervous system. While oxidative and nitrosative stress (RONS), resulting from an imbalance of reactive oxygen species (ROS) and reactive nitrogen species (RNS), is known to significantly contribute to the progression^14–17^, the respective contributions of ROS and RNS to αSyn propagation, and their relative pathogenic importance, remain largely unresolved. This fundamental gap in our understanding stems from the highly reactive and intertwined nature of these species in biological milieus^16–18^, coupled with a lack of tools for their selective modulation in relevant disease models.

To overcome this challenge and create precise tools to dissect the RONS network, we pursued a strategy centered on designing nanomaterials with tailored redox-modulating enzyme-like activities^19–21^. While emerging studies have identified select nanozymes with broad-spectrum antioxidant properties capable of scavenging both ROS and RNS^22–27^, achieving concurrent ROS elimination alongside controlled RNS generation remains a significant challenge^28,29^. Our primary objective was thus to develop two classes of nanozymes with opposing functionalities: one capable of broadly scavenging a wide range of RONS, and a counterpart designed to create a specific nitrosative stress environment by catalytically promoting nitric oxide (NO**•**) production, while simultaneously scavenging key ROS such as superoxide (O□□•), hydrogen peroxide (H□O□), and the hydroxyl radical (•OH). This strategy was informed by our prior research establishing the RNS-activated poly(ADP-ribose) (PAR) pathway as a potent driver of αSyn neurodegeneration^6^ and by compelling literature linking copper (Cu) exposure to α-synucleinopathies^30^ and its ability to modulate redox pathways^31–33^. The overarching goal was thus to leverage these specifically engineered nanozymes to dissect the respective roles and relative pathogenic importance of RNS and ROS in driving αSyn propagation.

Here, we report the successful realization of this strategy. Through a material-centric screening and design process, we developed two lead nano-probes —a pristine iridium (Ir) nanozyme that acts as a potent broad-spectrum RONS scavenger, and a copper-tuned iridium (Ir□Cu) nanozyme engineered to maintain superior ROS-scavenging capabilities while paradoxically amplifying RNS. This unique pair of nanozymes provided a platform to interrogate the differential roles of RNS and ROS *in vivo*. Employing these tools, we found that the RNS-amplifying Ir□Cu nanozyme dramatically exacerbated αSyn pathology despite being a more potent ROS scavenger. Through a multifaceted investigation encompassing detailed pathological, pharmacological, and transcriptomic analyses, our results point to the pathogenic primacy of the RNS pathway induced by Ir_3_Cu. Based on this insight, we then demonstrate that Ir nanozyme capable of scavenging both ROS and RNS provides neuroprotection and rescues functional deficits across multiple mouse models of α-synucleinopathy, including those that recapitulate brain-first, body-first, and AD/PD co-pathologies. This work, within our experimental models, suggests that RNS plays a predominant role in driving αSyn propagation and that the combination of RONS scavenging is a potential powerful therapeutic strategy for these devastating disorders.

## Rational design and cellular validation of iridium-copper nanozymes with opposing redox-modulating functions

To develop nanozyme tools capable of scavenging RONS in αSyn pathology propagation, we first performed a systematic screening of eight mono-metal nanozymes (Ru, Rh, Pd, Ag, Au, Pt, Ir, Cu) and carbon dots (CDs). A series of characterization revealed that while most candidates exhibited multiple enzyme-like activities (e.g., peroxidase-, superoxide dismutase-, and catalase-like) for scavenging various ROS including •OH, iridium (Ir) displayed superior broad-spectrum efficiency (Fig. 1a,b). However, these materials showed divergent behaviors in regulating RNS (Fig. 1c). Inspired by the classic reaction of copper-catalyzed S-nitrosothiol compounds to release NO• in biological systems^34–39^, we observed that while most of our tested nanozymes scavenged NO•, the copper (Cu) nanozyme uniquely promoted its generation through the catalytic decomposition of S-nitroso-N-acetylpenicillamine (SNAP)^40^ (Fig. 1c).

**Fig. 1.**
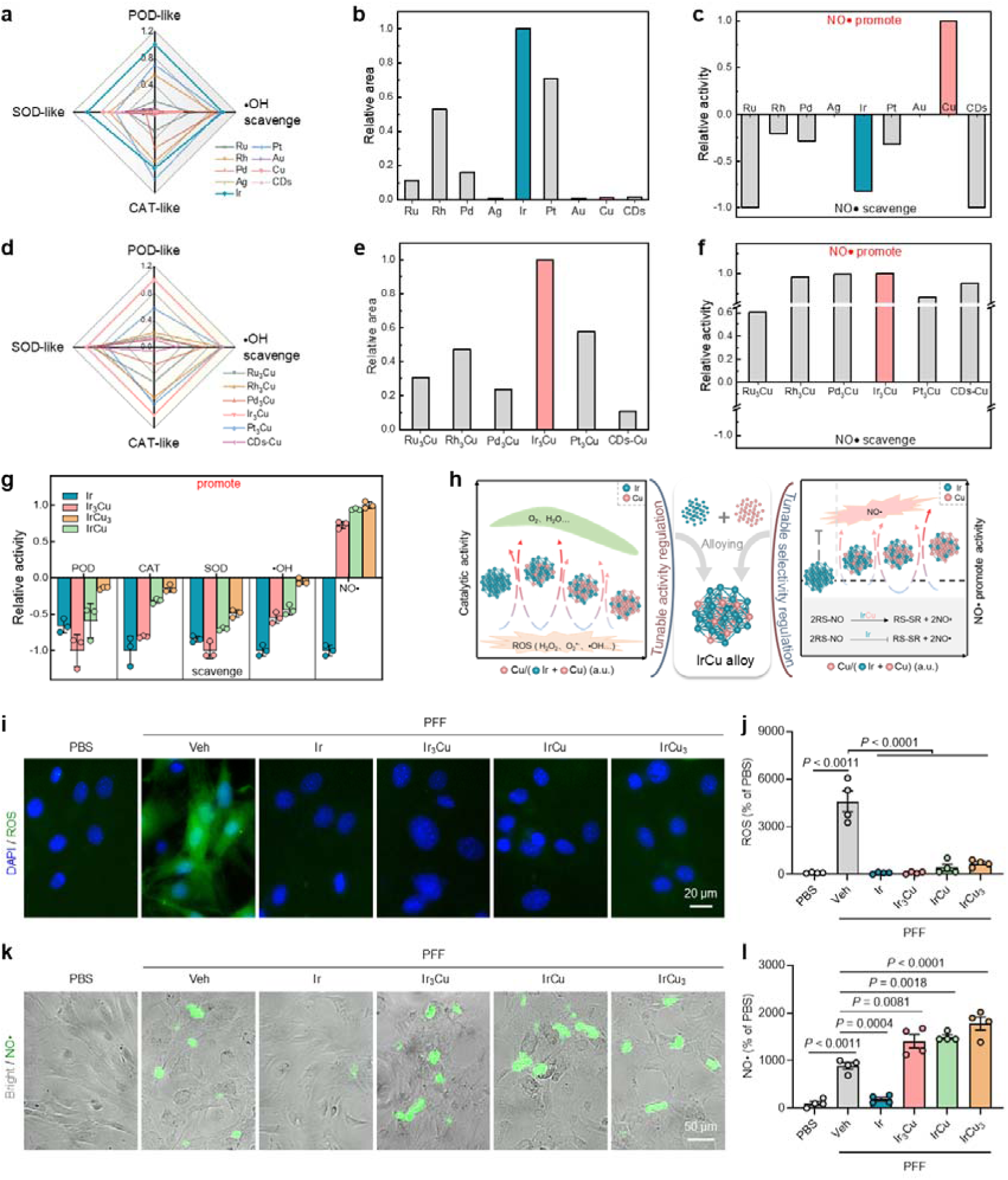
Rational design and cellular validation of iridium-copper nanozymes with opposing redox-modulating functions. **a**, A comparison of the relative activities of ROS-related enzymes (POD-, CAT-, SOD-like and •OH scavenging capabilities) of various mono metal NPs and CDs. **b**, A comparison of the overall relative anti-ROS capabilities of various mono metal NPs and CDs. The relative areas enclosed by various mono metal NPs and CDs were calculated through (**a**). **c**, A comparison of the relative activities of various mono metal NPs and CDs catalysts in promoting or scavenging NO• production. **d**, A comparison of the relative activities of ROS-related enzymes (POD-, CAT-, SOD-like and •OH scavenging capabilities) of various Cu based nanozymes (bimetallic alloy and CDs-Cu). **e**, A comparison of the overall relative anti-reactive oxygen species capabilities of various Cu based nanozymes. The relative areas enclosed by various Cu based nanozymes were calculated through (**d**). **f**, A comparison of the relative activities of various Cu based nanozymes in promoting or scavenging NO• production. **g**, A comparison of the relative activities of Ir and Ir-Cu nanozymes with different Ir/Cu ratio in scavenging or promoting ROS-related enzymes (POD, CAT, SOD and •OH scavenging capabilities) and NO• production. Data are presented as mean ± SEM, *n* = 3. **h**, Schematic illustration of Cu tuning the catalytic activity of Ir to scavenge ROS and selectivity towards NO• (promotion or scavenging). By introducing Cu atoms, the alloying of Ir-Cu regulates the scavenging activity of ROS and the selectivity in the switching of Ir from scavenging to promoting the production of NO•. For (**a–g**) the relative activity value or area of the material with the highest enzyme activity or the largest enclosed area is defined as 1, and the relative activity value or area of other materials is the ratio of that material. For (**c**,**f**,**g**) the positive/negative sign before the relative activity value represents promotion and scavenging respectively. **i–l**, Mouse primary microglia were treated with vehicle (Veh) or various nanozymes (Ir, Ir_3_Cu, IrCu, or IrCu_3_) in the presence of either αSyn PFF or PBS. **i,** Representative images and (**j**) corresponding quantification of ROS signals in primary microglia, with ROS coverage normalized to the PBS group. Nuclei are counterstained with DAPI (blue). **k,** Representative images and (**l**) corresponding quantification of NO• signals in primary microglia, with the percentage of NO•-positive cells normalized to the PBS group. *n* = 4. Data are presented as mean ± SEM. *P* < 0.05 by one-way ANOVA with Tukey’s multiple comparisons test.

We find that while Ir is a superior RONS scavenger, Cu is a specific RNS promoter. As such, we hypothesized that Cu could be used as a dopant to impart RNS-promoting capabilities onto other potent ROS-scavenging nanozymes. To test this principle, we synthesized a series of bimetallic or hybrid nanozymes by combining Cu with various ROS-scavenging metals or carbon dots (CuM, where M= Ru, Rh, Pd, Ir, Pt, CDs). As anticipated, the Cu-containing nanozymes not only retained the inherited ROS-scavenging activities of their parent materials but universally demonstrated a robust catalytic capacity to promote NO• production (Fig. 1d–f). This confirmed our hypothesis and established “copper-tuning” as a generalizable strategy to create nanozymes capable of scavenging RONS and other nanozymes capable of promoting RNS.

To identify the optimal system for our study, we then systematically investigated nanozymes with varying Ir:Cu stoichiometric ratios (Ir, Ir□Cu, IrCu, IrCu□). This direct comparison established a clear composition-activity profile: ROS scavenging efficiency was highest with high Ir content (following the hierarchy of Ir□Cu > IrCu > IrCu□), while robust NO• generation required the incorporation of Cu (Fig. 1g,h and Extended Data Figs. 1 and 2). This screening identified Ir and Ir□Cu as the two most compelling candidates, as they exhibited comparable, top-tier ROS-scavenging capabilities, yet possessed diametrically opposite functions in NO• regulation. We therefore proceeded with their in-depth physicochemical characterization. The nanozymes are ultrasmall and coated with polyvinylpyrrolidone (PVP) for excellent water solubility and biocompatibility (Extended Data Figs. 1 and 2). Detailed characterization (HRTEM, XRD, STEM-EDS) confirmed the formation of homogeneous alloy nanoparticles with a face-centered cubic structure.

Having characterized the intrinsic catalytic properties of our lead nanozymes, we next sought to validate their redox-modulating functions in a relevant cellular context. We used primary microglia stimulated with αSyn PFF to mimic the neuroinflammatory environment of α-synucleinopathy^41,42^. As anticipated from our cell-free assays, both Ir and Ir□Cu nanozymes potently suppressed the αSyn PFF-induced ROS surge in these cells (Fig. 1i,j). In contrast, while the Ir nanozyme also scavenged the αSyn PFF-induced NO• production (Fig. 1k,l), the Ir□Cu nanozyme dramatically amplified the NO• signal to levels far exceeding those induced by αSyn PFF alone while potently reducing ROS. In brief, this material-centric design, screening, characterization, and cellular validation effort yielded an ideal pair of nano-probes—Ir (a potent RONS scavenger) and Ir□Cu (a potent ROS scavenger that also acts as an RNS amplifier).

## ROS-scavenging and NO• promoting Ir□Cu nanozyme exacerbates αSyn pathology

Having established the distinct and opposing redox-modulating properties of our Ir and Ir□Cu nanozymes *in vitro*, we next investigated their effects on αSyn pathology *in vivo*. We employed an established mouse model where pathological αSyn propagation is initiated by a single stereotactic injection of αSyn PFF into the striatum^6–13^, followed by weekly intranasal administration of vehicle (Veh), Ir, or Ir□Cu nanozymes (Fig. 2a). As expected, PFF injection induced a substantial and widespread dissemination of pathological, phosphorylated Ser129 (pS129)-positive αSyn inclusions throughout interconnected brain regions (Fig. 2b–d). The two nanozymes produced opposite effects on this pathology. While intranasal administration of the Ir nanozyme attenuated αSyn pathology, the Ir□Cu nanozyme treatment, in contrast, failed to suppress it and instead significantly exacerbated the pS129 burden to levels beyond that of the PFF-only group. This exacerbation by Ir□Cu and suppression by Ir was consistently observed across multiple key brain regions, including the prefrontal cortex (PFC), cortex (CTX), amygdala (AMG), and substantia nigra pars compacta (SNc) (Fig. 2b–d), and was accompanied by a corresponding increase in Ir□Cu-driven microglial activation (Extended Data Fig. 3a–c).

**Fig. 2.**
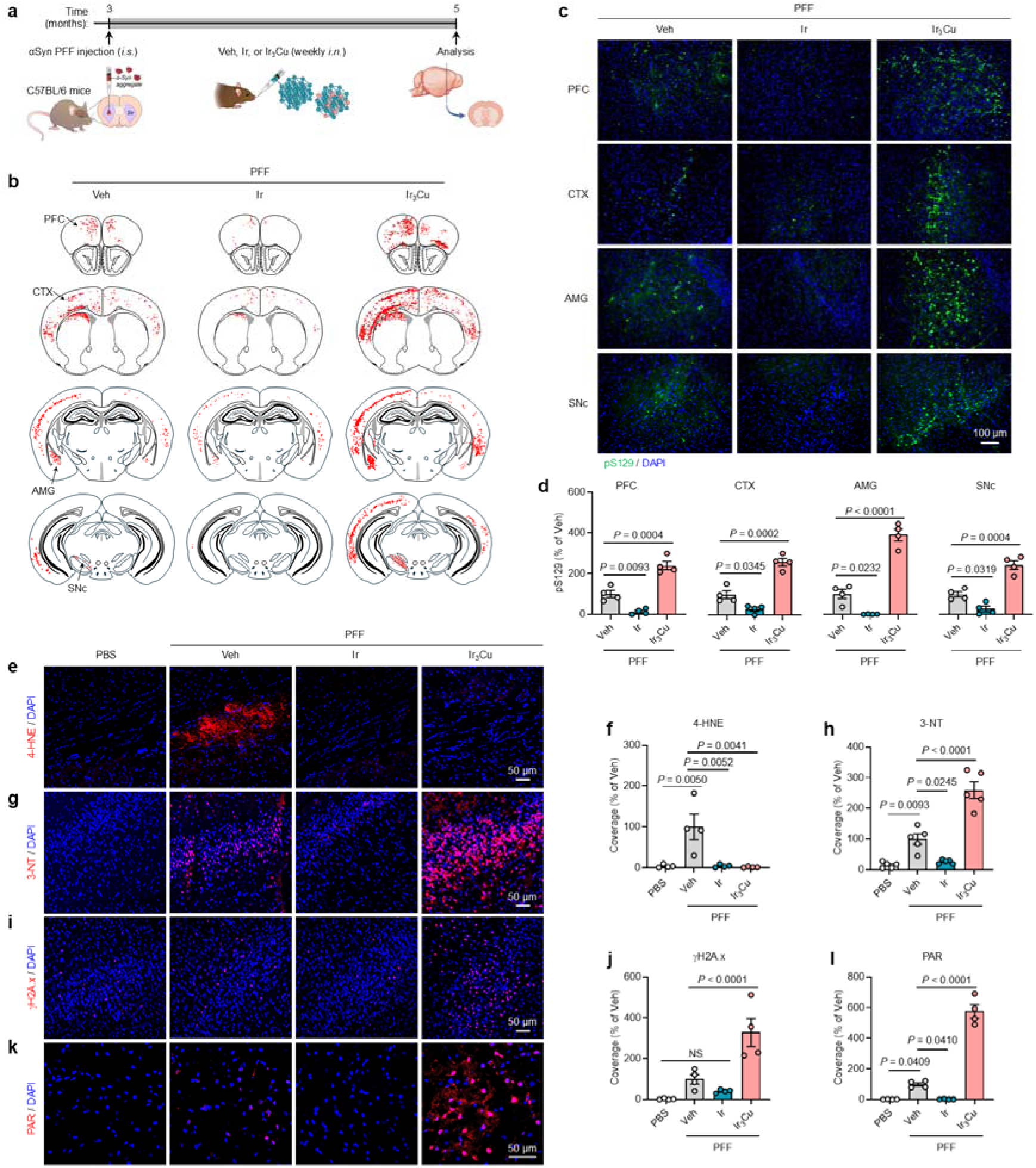
ROS-scavenging Ir□Cu nanozyme paradoxically exacerbates pathology, revealing RNS dominance *in vivo*. **a**, Schematic of the experimental workflow: C57BL/6 wide-type mice received intrastriatal (*i.s.*) injection of αSyn PFF, followed by weekly intranasal (*i.n.*) administration of vehicle (Veh) or nanozymes (Ir or Ir_3_Cu). Brains were collected 2 months after αSyn PFF injection. **b**, Dot map representation of the distribution of pS129-positive αSyn pathology in mice. **c**, Representative fluorescent images showing pS129-positive αSyn pathology (green) in prefrontal cortex (PFC), cortex (CTX), amygdala (AMG), and substantia nigra pars compacta (SNc). **d**, Corresponding quantification of the coverage of pS129-positive αSyn pathology normalized to the Veh group. **e-l**, Representative images and corresponding quantification of (**e**,**f)** 4-hydroxynonenal (4-HNE), (**g**,**h**) 3-nitrotyrosine (3-NT), (**i**,**j**) γH2A.x, and (**k**,**l**) poly(ADP-ribose) (PAR), with coverage normalized to Veh group. Nuclei are stained with DAPI (blue). Data are presented as mean ± SEM. *n* = 4–5 mice per group. *P* < 0.05 by one-way ANOVA with Tukey’s multiple comparisons test. NS, not significant.

To dissect the underlying mechanisms driving these opposing pathological outcomes, we quantified key markers of oxidative/nitrosative stress and downstream DNA damage. PFF injections led to a pronounced increase in 4-hydroxynonenal (4-HNE), a product of ROS-driven lipid peroxidation^14,17,18^, and 3-nitrotyrosine (3-NT), a specific marker for RNS peroxynitrite (ONOO□)-mediated protein nitration^14–17,43^ (Fig. 2e–h). Consistent with their ROS-scavenging capabilities, both Ir and Ir□Cu nanozymes significantly reduced 4-HNE levels *in vivo* (Fig. 2e,f). Their effects on 3-NT, however, were significantly different: Ir treatment suppressed 3-NT levels, whereas Ir□Cu treatment caused their elevation beyond PFF-only controls (Fig. 2g,h). This suggests that the exacerbated αSyn pathology tracks with the amplified nitrosative stress, not bulk ROS levels. Further supporting this, we found that the downstream DNA damage response pathway, previously linked to RNS^6^, followed the same pattern. The RNS-amplifying Ir□Cu nanozyme caused an accumulation of the DNA damage marker γH2A.x and the Poly(ADP-ribose) polymerase-1 (PARP-1) activation product poly(ADP-ribose) (PAR), while the neuroprotective Ir nanozyme suppressed both (Fig. 2i–l).

In brief, these results provide evidence of a strong *in vivo* correlation between exacerbated αSyn pathology and the RNS-PARP-PAR cascade, but not with markers of ROS. To determine if this RNS-associated pathology is indeed causal, we next investigated whether pharmacological inhibition of RNS synthesis could prevent the pathology driven by the Ir□Cu nanozyme.

## Pharmacological and transcriptomic evidence for the causal role of Ir_3_Cu-related RNS in **α**-synuclein pathology

To determine whether Ir_3_Cu-promoted NO• generation is causally involved in αSyn pathology, we administered the broad-spectrum nitric oxide synthase (NOS) inhibitor, *N*^G^-nitro-L-arginine methyl ester (L-NAME)^44–48^, to PFF-injected mice co-treated with the Ir□Cu nanozyme (hereafter referred to as Ir□Cu-LN) (Fig. 3a). L-NAME administration abrogated the exacerbation of pS129-positive αSyn pathology induced by Ir□Cu, reducing the pathological burden to PFF-only baseline levels (Fig. 3b,c). This rescue of the pathological phenotype was accompanied by a corresponding reduction in microglial activation (Extended Data Fig. 3d–f). Consistent with the inhibition of NO• production, L-NAME also suppressed the downstream molecular cascade. In the Ir□Cu-LN group, the elevated levels of the protein nitration marker 3-NT, the DNA double-strand break marker γH2A.x, and the PARP activation product PAR were all decreased compared to the Ir□Cu-only group (Fig. 3d–i). These findings provide evidence linking RNS overproduction by Ir□Cu to αSyn pathology and its downstream molecular effectors.

**Fig. 3.**
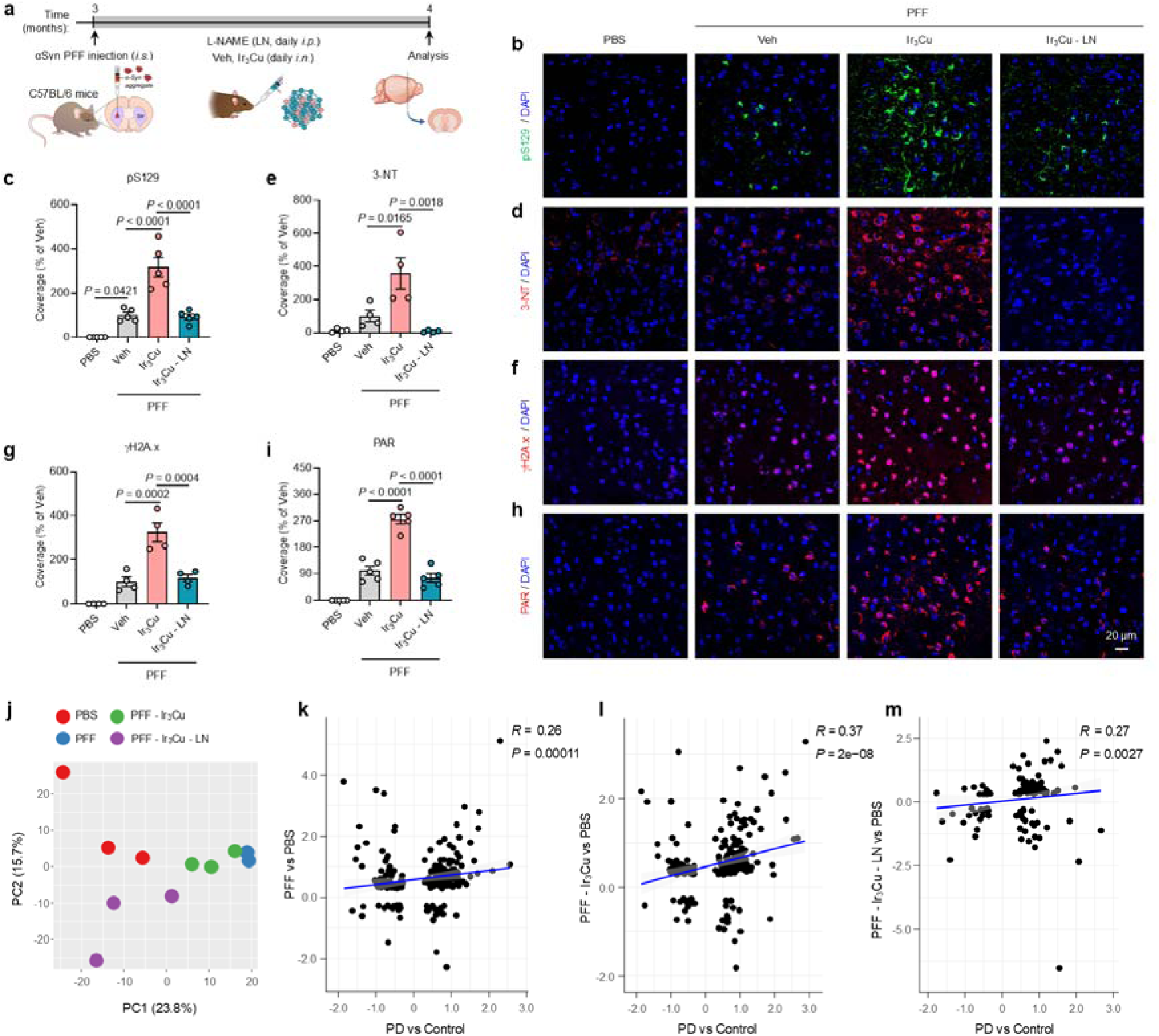
Pharmacological and transcriptomic evidence for the causal role of Ir_3_Cu-related RNS in α-synuclein pathology. **a**, Schematic of the *in vivo* experimental design showing stereotactic injection of αSyn PFF into the striatum of C57BL/6 mice, followed by *i.n.* administration of Veh, or Ir□Cu nanozyme combined with the intraperitoneal (*i.p.*) administration of the nitric oxide synthase inhibitor L-NAME (LN). Brains were collected 1 month after αSyn PFF injection. **b–i**, Representative images and corresponding quantification of (**b**,**c**) pS129 (green), (**d**,**e**) 3-NT (red), (**f**,**g**) γH2A.x (red), and (**h**,**i**) PAR (red) in the cortex region, with coverage normalized to Veh group. Nuclei are counterstained with DAPI (blue). Data are presented as mean ± SEM. *n* = 4–5 mice per group. *P* < 0.05 by one-way ANOVA with Tukey’s multiple comparisons test. **j–m,** C57BL/6 WT mice received αSyn PFF and treatment, cortex containing anterior cingulate cortex (ACC) region was isolated for total RNA extraction and RNA-seq. **j**, Principal component analysis (PCA) of transcriptional profiles based on the top 1,000 most variable genes. **k–m**, Scatterplots showing the correlation between transcriptional changes in treated mice (**k**, PFF vs PBS; **l**, PFF-Ir_3_Cu vs PBS, **m**, PFF-Ir_3_Cu-LN vs PBS) and those observed in the ACC of PD patients (PD vs Control).

To further investigate the molecular impact of Ir□Cu-promoted NO• signaling, we performed RNA-sequencing on cortical tissue to evaluate the transcriptomic responses induced by PFF, PFF-Ir_3_Cu, and PFF-Ir_3_Cu-LN inoculation in wide-type (WT) mice, compared to PBS controls. Principal component analysis (PCA) of the top 1,000 most variable genes revealed a clear separation between all groups (Fig. 3j); notably, the PFF-Ir□Cu samples formed a distinct cluster that was shifted back towards the PFF and PBS controls by L-NAME treatment (Fig. 3j). Differential gene expression (DEG) analysis identified the largest number of differentially expressed genes (DEGs; *P_adj_* < 0.05) in the PFF-Ir_3_Cu group (*n* = 391), followed by PFF (*n* = 368), with the fewest DEGs observed in the PFF-Ir_3_Cu-LN condition (*n* = 134) (Extended Data Fig. 4a,b and Supplementary Tables 1–3). Comparison of DEGs across conditions showed both unique and shared responses (Extended Data Fig. 4a). Among the 391 DEGs in the PFF-Ir□Cu group, the majority (236) were unique to this condition, while only a small fraction overlapped with the PFF (*n* = 84) or PFF-Ir□Cu-LN (*n* = 30) groups (Extended Data Fig. 4a). This limited gene overlap provides quantitative evidence that L-NAME treatment substantially reverses the specific transcriptional program driven by Ir□Cu-induced RNS. All conditions showed more upregulated than downregulated genes, with immune-related transcripts such as *C3*, *C4b*, and *Cxcl12* being prominently induced in the PFF-Ir□Cu group and dampened in the PFF-Ir□Cu-LN group, while downregulated genes like *Egr2* and *Sox1* were fewer and more condition-specific (Extended Data Fig. 4b).

Gene set enrichment analysis (GSEA) further revealed the greatest number of dysregulated gene sets in the PFF-Ir_3_Cu condition (*n* = 223), followed by PFF-Ir_3_Cu-LN (*n* = 101), and the fewest in PFF (*n* = 86) (Extended Data Fig. 4c, Supplementary Figs. 1–3 and Supplementary Tables 1–3). In the PFF-Ir_3_Cu group, most gene sets (193) were upregulated and related to immune processes, while the 30 downregulated sets were primarily involved in neuronal pathways (Supplementary Fig. 1 and Supplementary Table 1). Similarly, in PFF-Ir_3_Cu-LN, nearly all gene sets (*n* = 100) were upregulated and immune-related, and only one was downregulated and involved in forebrain regionalization (Supplementary Fig. 2 and Supplementary Table 2). In contrast, the PFF condition showed a dominance of downregulated gene sets (63), primarily associated with neuronal functions, while only 23 gene sets were upregulated, mainly related to the immune system (Supplementary Fig. 3 and Supplementary Table 3). The smaller set of upregulated immune-related pathways in PFF-Ir_3_Cu-LN relative to PFF-Ir_3_Cu indicates that L-NAME mitigates RNS-driven inflammatory signaling initiated by Ir_3_Cu.

We further compared gene expression changes in each mouse condition with transcriptomic profiles from the anterior cingulate cortex of PD patients (Fig. 3k–m). While PFF-treated mice showed a modest but significant correlation with human PD profile (Spearman correlation, *R* = 0.26, *P* = 1.7 × 10□□; Fig. 3k), the PFF-Ir_3_Cu mice exhibited a much stronger correlation (*R* = 0.37, *P* = 2.0 × 10□□; Fig. 3l). In contrast, this enhanced correlation was abrogated in the PFF-Ir_3_Cu-LN group (*R* = 0.27, *P* = 0.0027; Fig. 3m). Thus, at the whole-transcriptome level, RNS amplification by Ir□Cu pushes the PFF model’s gene expression profile closer to that of human PD, an effect that is abrogated by RNS inhibition.

Taken together, the pharmacological and transcriptomic data provide evidence for RNS being a primary pathogenic driver in α-synucleinopathies. These data demonstrate that in this αSyn PFF-induced PD mice model, the Ir□Cu nanozyme induces an overabundance of RNS, which in turn drives a DNA damage-PAR cascade and a PD-like transcriptomic signature, ultimately exacerbating αSyn pathology. This pathogenic pathway is summarized in a schematic model (Extended Data Fig. 5).

## Intranasal delivery of the RONS-scavenging Ir nanozyme provides neuroprotection and rescues behavioral deficits in a brain-first PD model

The pathogenic potential of RNS, revealed by the paradoxical effects of the Ir□Cu nanozyme, provided a rationale for testing the therapeutic efficacy of a nanozyme designed to scavenge RNS. We therefore evaluated our broad-spectrum RONS-scavenging Ir nanozyme *in vitro* and *in vivo*.

We first confirmed its mechanism of action at the cellular level. We employed a human embryonic kidney cell line (HEK293)^49,50^ engineered to express a familial PD-associated αSyn mutant (A53T) fused to yellow fluorescent protein (YFP), hereafter referred to as HEK-A53T-YFP. Upon stimulation with αSyn PFF, these cells developed intracellular αSyn aggregates, as indicated by enhanced punctate YFP fluorescence (Extended Data Fig. 6a,b). Confocal imaging and line-scan analysis confirmed efficient uptake of Cy5.5-labeled Ir nanozyme by HEK-A53T-YFP cells (Extended Data Fig. 6c). When αSyn PFF was added 3 hours prior to Ir nanozyme treatment, subsequent αSyn aggregation was partially suppressed (Extended Data Fig. 6d,e), Moreover, co-treatment with Ir nanozyme and αSyn PFF significantly reduced the proportion of cells exhibiting pathological punctate YFP signals (Extended Data Fig. 6f,g), indicating that Ir nanozyme treatment effectively inhibits αSyn seeding and aggregation. In primary neurons exposed to αSyn PFF, the Ir nanozyme also led to a marked reduction in pS129-positive αSyn aggregates (Extended Data Fig. 6h,i). These *in vitro* results demonstrate that the Ir nanozyme can directly inhibit the core pathogenic process of seeded αSyn aggregation in neuronal cells and human cells.

To determine if this direct anti-aggregation activity translates to *in vivo* efficacy, we first found that intranasal (*i.n.*) administration is a highly efficient method for delivering nanozymes to the brain, leading to substantially higher brain enrichment compared to the intravenous (*i.v.*) route (Extended Data Fig. 7a–c). We then tested the therapeutic effects of both delivery routes in our striatal PFF injection mouse model (Fig. 4a). Weekly treatment with the Ir nanozyme via either route significantly attenuated the accumulation of pS129-positive αSyn aggregates in the cortex compared to vehicle-treated controls. Notably, intranasal delivery exhibited a significantly superior inhibitory effect on αSyn pathology compared to the intravenous route (Fig. 4b,c and Extended Data Fig. 7d,e). Furthermore, PFF-induced microglial activation (IBA1-positive cells) was also suppressed by Ir nanozyme treatment; in this case, both *i.n.* and *i.v.* administration showed comparable efficacy in reducing neuroinflammation (Fig. 4b,d). Given its superior brain-targeting efficiency and greater anti-pathology effect, the intranasal route was selected for all subsequent therapeutic assessments.

**Fig. 4.**
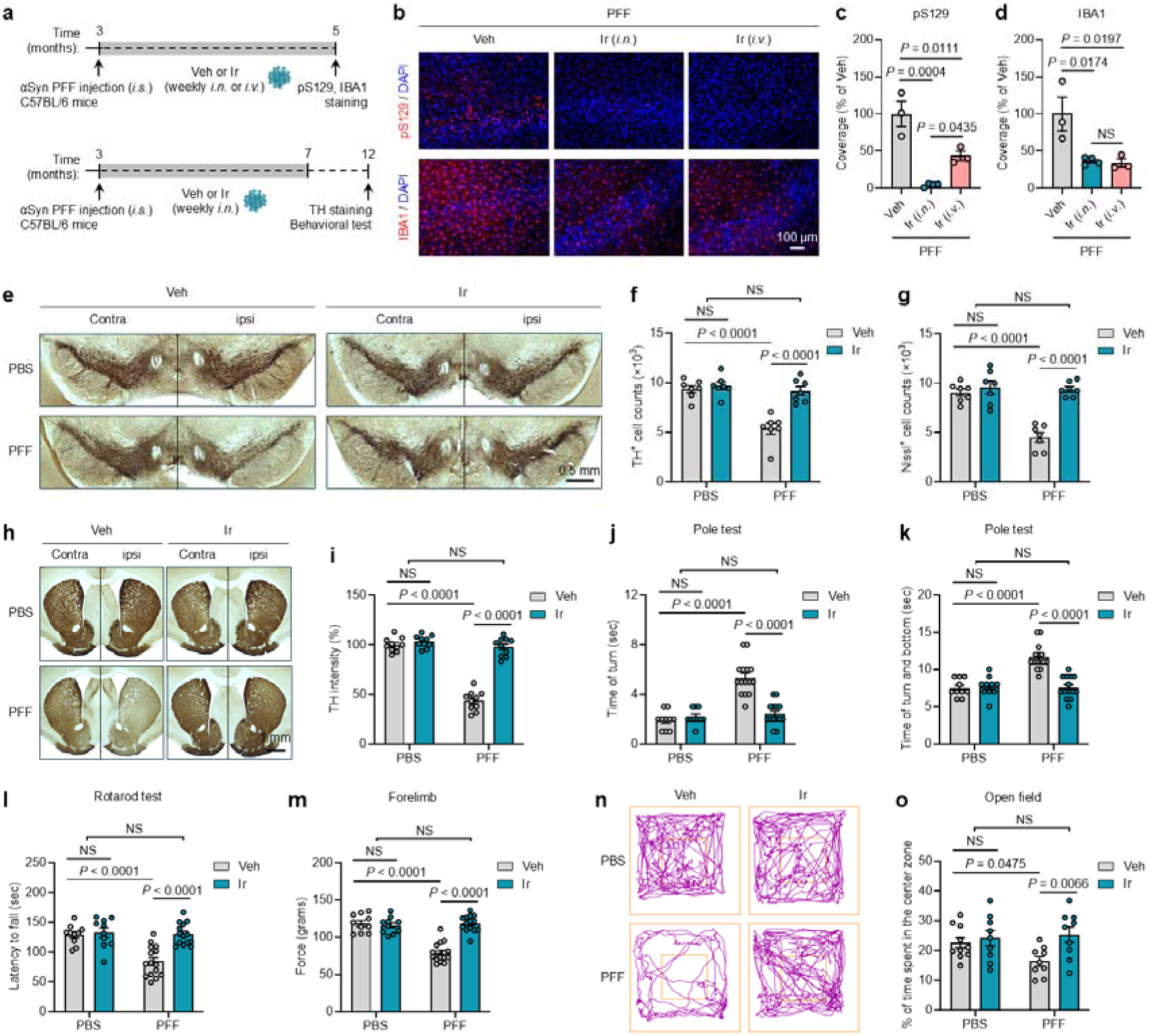
Intranasal delivery of the RONS-scavenging Ir nanozyme provides neuroprotection and rescues behavioral deficits in a brain-first PD model. **a**, Schematic showing C57BL/6 WT mice received *i.s.* αSyn PFF injection, followed by weekly *i.n.* or *i.v.* administration of Ir nanozyme. Brains were collected 2 months post-PFF injection to assess αSyn pathology and neuroinflammation. For tyrosine hydroxylase (TH) staining and behavioral testing, C57BL/6 WT mice received *i.s.* αSyn PFF or PBS and weekly *i.n.* Ir nanozyme or vehicle for 4 months; assessments were performed 9 months after PFF injection. **b**, Representative images showing pS129-positive αSyn pathology (red) and IBA1-positive microglia (red) in the cortex of PFF-injected mice treated with Ir nanozyme *i.n.* or *i.v.* route. Nuclei are counterstained with DAPI (blue). **c**,**d**, Quantification of (**c**) pS129 and (**d**) IBA1, with coverage normalized to the Veh group. *n* = 3**–**4 mice per group. **e**, TH and Nissil staining in the substantia nigra. **f**,**g**, Stereological quantification of (**f**) TH-positive cells and (**g**) Nissil-positive cells in the substantia nigra. *n* = 7 mice per group. **h**, Representative images of TH and Nissil staining in the striatum. **i**, Quantification of TH fiber density in the striatum. *n* = 10 mice per group. **j–o**, Behavioral assessment of Veh- or Ir- treated mice at 9 months after *i.s.* αSyn PFF or PBS injection. Results of mice on the (**j**,**k**) pole test: time to turn and total time to descend, (**l**) rotarod test: latency to fall, (**m**) forelimb grip strength test: peak force, and (**n**,**o**) open field test: percent time spent in the center zone are shown. **j–m**, *n* = 10 mice for PBS-Veh group, *n* = 15 mice for PFF-Veh group, *n* = 10 mice for PBS-Ir group, *n* = 14 mice for PFF-Ir group. For **o**, *n* = 10 mice for PBS-Veh group, *n* = 9 mice for PFF-Veh group, PBS-Ir group, or PFF-Ir group. Data are presented as mean ± SEM. *P* < 0.05 by one-way ANOVA with Tukey’s multiple comparisons test for **c** and **d**. *P* < 0.05 by two-way ANOVA with Tukey’s multiple comparisons test for **f**, **g**, **i–m**, and **o**.

We next assessed whether this suppression of pathology by intranasal Ir nanozyme treatment translated into dopaminergic neuroprotection and a corresponding rescue of functional deficits. Nine months after PFF injection, vehicle-treated mice exhibited neurodegeneration of the nigrostriatal pathway (Fig.4a, e–i). Stereological counts in the substantia nigra pars compacta (SNc) revealed an approximately 50% loss of both tyrosine hydroxylase (TH)-positive dopaminergic neurons and the total Nissl-positive neuronal population (Fig. 4e–g). This loss of neuronal cell bodies was accompanied by a severe reduction of their striatal terminals, as measured by TH fiber density (Fig. 4h,i). In contrast, intranasal Ir nanozyme treatment effectively prevented this neuronal loss in the SNc (Fig. 4e–g), and fiber loss (Fig. 4h,i).

This neuroprotection at the cellular and circuit level translated into a rescue of functional abilities across a battery of behavioral tests. In the pole test, PFF-injected mice showed significantly impaired motor coordination, taking longer to turn and descend the pole; Ir nanozyme treatment prevented this deficit, with performance statistically indistinguishable from PBS-injected controls (Fig. 4j,k). Similarly, in the rotarod test, the reduced latency to fall caused by PFF injection was rescued by Ir nanozyme treatment (Fig. 4l). The PFF-induced reduction in muscular strength was also reversed, as shown by the forelimb grip strength test, where the Ir-treated group performed at control levels (Fig. 4m). Furthermore, we assessed anxiety-like and exploratory behaviors using the open field test. PFF-injected mice spent significantly less time in the center of the arena, a deficit that was normalized by Ir nanozyme administration (Fig. 4n,o). These behavioral analyses demonstrated that Ir nanozyme treatment led to prevention of both motor and non-motor functions in this model of PD.

Interestingly, PFF-injected mice exhibited splenomegaly, indicative of peripheral immune activation, and this was also alleviated by intranasal Ir nanozyme treatment (Extended Data Fig. 7f–h). Taken together, these findings demonstrate that the RONS-scavenging Ir nanozyme, delivered non-invasively, not only halts central disease progression by directly inhibiting αSyn pathology and protecting the dopaminergic system but also mitigates associated peripheral inflammation, leading to a systematic rescue of functional deficits.

## Oral Ir nanozyme treatment blocks gut-to-brain **α**-synuclein propagation in a peripheral-origin PD model

Building on the therapeutic efficacy demonstrated in the brain-first model, we next sought to determine whether the Ir nanozyme could inhibit αSyn pathology in a “body-first” disease context. This line of inquiry was motivated by the Braak hypothesis and extensive clinical evidence suggesting that αSyn pathology can originate in the peripheral nervous system, particularly the gut, and propagate to the brain via the vagus nerve^3,51–53^. To model this, we injected αSyn PFFs into the gastric wall of A53T αSyn transgenic mice^54^, followed by weekly oral gavage administration of the Ir nanozyme or vehicle for two months (Fig. 5a).

**Fig. 5.**
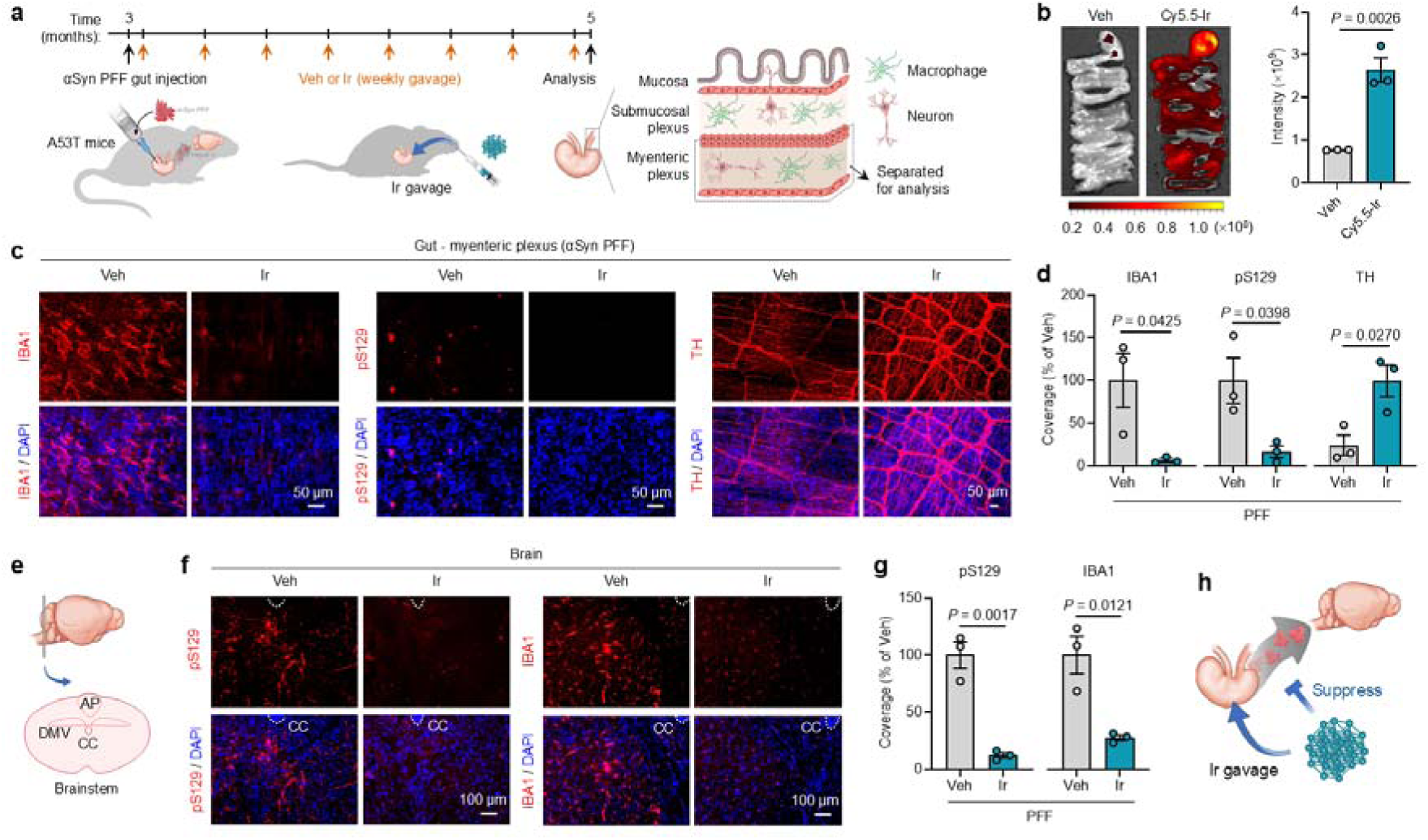
Intranasal delivery of the RONS-scavenging Ir nanozyme mitigates converging co-pathologies and rescues neuronal loss in a model of Alzheimer’s disease with Lewy body pathology. **a**, Schematic of the experimental workflow: 9-month-old 5XFAD mice received intra-hippocampal (*i.h.*) injection of αSyn PFF, followed by weekly *i.n.* administration of Ir nanozyme or vehicle. Brains were collected 3 months post-PFF injection for analysis. **b–d**, Representative images and corresponding quantification of (**c**) pS129-postive p-αSyn (green) and (**d**) AT8-positive p-tau (red) pathology in the cortex. *n* = 3 mice per group. **e**, Dot map representation of the distribution of pS129-positive αSyn pathology in the mice brain. **f**, Representative images showing pS129-positive p-αSyn pathology, Aβ_42_-positive amyloid deposition, AT8-positive p-tau pathology, and IBA1-positive microglia in the cortical brain region of aSyn PFF-injected 5XFAD mice treated with Ir nanozyme or vehicle. **g**, Corresponding quantification of cortical pathology pS129, Aβ_42_, AT8, IBA1, with coverage normalized to the Veh group. *n* = 4 mice per group. **h**, Immunofluorescence staining of NeuN-positive neurons in the hippocampus, including cornu ammonis 1 (CA1) and dentate gyrus (DG), and **i**, corresponding quantification of NeuN-positive cells per field of view (FOV) in each region. *n* = 4 mice per group. Data are presented as mean ± SEM. *P* < 0.05 by unpaired two-tailed Student’s *t*-test.

We first confirmed that orally administered Ir nanozymes can be effectively delivered and retained within the gastrointestinal tract, as shown by *in vivo* imaging of Cy5.5-labeled Ir nanozymes (Fig. 5b). At the site of pathological initiation, the myenteric plexus (Fig. 5a), Ir nanozyme treatment demonstrated local efficacy. It significantly suppressed PFF-induced the activation of enteric macrophages (IBA1-positive cells) (Fig. 5c,d), markedly decreased the accumulation of pathological pS129-positive αSyn aggregates (Fig. 5c,d), and preserved the integrity of dopaminergic fibers, as evidenced by restored TH expression in the gut (Fig. 5c,d).

Having established the potent local effects of the Ir nanozyme in the gut, we next investigated the question of whether this peripheral intervention could block the trans-neuronal propagation of αSyn pathology to the central nervous system. We focused our analysis on the dorsal motor nucleus of the vagus (DMV) in the brainstem, a key initial site of CNS pathology in gut-first models (Fig. 5e). As expected, vehicle-treated mice showed significant accumulation of pS129-positive αSyn and associated neuroinflammation (IBA1-positive microglia) in the brainstem. Oral treatment with the Ir nanozyme prevented the appearance of this ascending pathology, with both pS129 and IBA1 signals being significantly reduced compared to controls (Fig. 5f,g).

These findings demonstrate that orally administered Ir nanozymes can effectively mitigate local inflammatory and pathological responses in the gut and block the trans-neuronal propagation of αSyn pathology to the brainstem (Fig. 5h). This result validates the therapeutic potential of the Ir nanozyme in a clinically relevant, peripheral-origin model of PD, significantly broadening its prospective application for treating different subtypes of α-synucleinopathies.

## Intranasal delivery of the RONS-scavenging Ir nanozyme mitigates converging co-pathologies and rescues neuronal loss in a model of Alzheimer’s disease with Lewy body pathology

A substantial portion of patients with AD also present with Lewy body (LB) co-pathology, and emerging evidence suggests that pre-existing amyloid-β (Aβ) plaques can potentiate the seeding and spreading of αSyn pathology^55^. To investigate whether our RONS-scavenging Ir nanozyme could intervene in this pathological crosstalk, we established a clinically relevant co-pathology model by injecting αSyn PFF into the hippocampus of 9-month-old 5XFAD mice, an established model of AD amyloidosis (Fig. 6a). This intervention successfully recapitulated a converging pathology state; PFF injection induced not only the accumulation of pS129-positive αSyn but also triggered the cross-seeding and hyperphosphorylation of endogenous tau (AT8-positive), a phenomenon not observed in PBS-injected 5XFAD controls (Fig. 6b–d and Extended Data Fig. 7i–k).

**Fig. 6.**
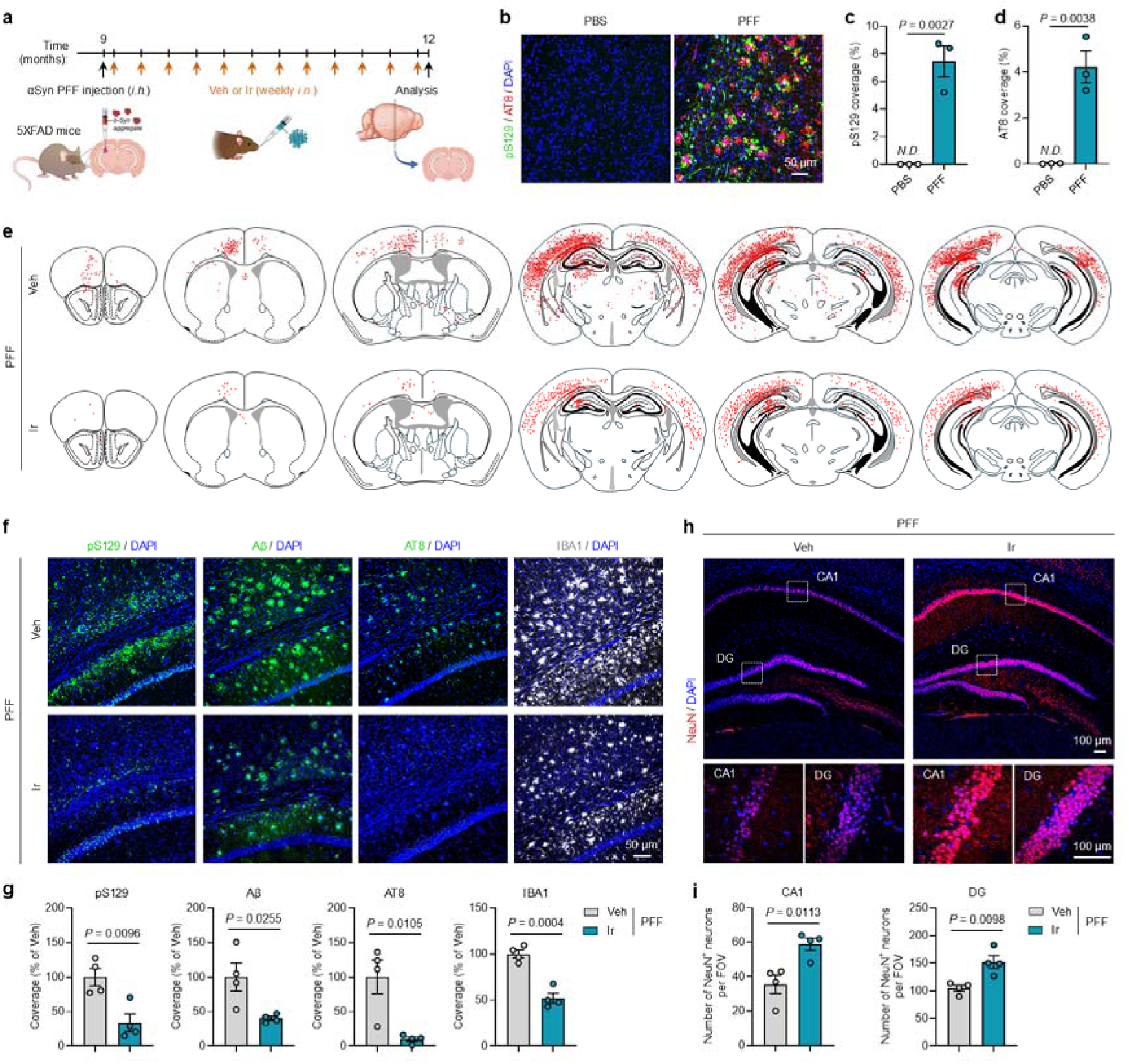
Ir nanozyme inhibits AD/PD co-pathology and rescues neuron loss in αSyn PFF-injected 5XFAD mice. **a**, Schematic of the experimental workflow: 9-month-old 5XFAD mice received intra-hippocampal (*i.h.*) injection of αSyn PFF, followed by weekly *i.n.* administration of Ir nanozyme or vehicle. Brains were collected 3 months post-PFF injection for analysis. **b**-**d**, Representative images and corresponding quantification of pS129-postive p-αSyn (green) (**c**) and AT8-positive p-tau (red) (**d**) pathology in the cortex. *n* = 3 mice per group. **e**, Dot map representation of the distribution of pS129-positive αSyn pathology in the mice brain. **f**, Representative images showing pS129-positive p-αSyn pathology, Aβ_42_-positive amyloid deposition, AT8-positive p-tau pathology, and IBA1-positive microglia in the cortical brain region of aSyn PFF-injected 5XFAD mice treated with Ir nanozyme or vehicle. **g**, Corresponding quantification of cortical pathology pS129, Aβ_42_, AT8, IBA1, with coverage normalized to the Veh group. *n* = 4 mice per group. **h**, Immunofluorescence staining of NeuN-positive neurons in the hippocampus, including cornu ammonis 1 (CA1) and dentate gyrus (DG), and **i**, corresponding quantification of NeuN-positive cells per field of view (FOV) in each region. *n* = 4 mice per group. Data are presented as mean ± SEM. *P* < 0.05 by unpaired two-tailed Student’s *t*-test.

Having established this co-pathology model, we assessed the therapeutic efficacy of a 3-month course of intranasal Ir nanozyme treatment. The Ir nanozyme demonstrated a multi-target effect, simultaneously mitigating all three core proteinopathies. Ir nanozyme treatment resulted in a marked reduction in pS129-positive αSyn aggregates, AT8-positive p-tau pathology, and the underlying Aβ_42_-positive plaque burden in the cortex compared to vehicle-treated controls (Fig. 6e–g). This broad anti-pathology effect was accompanied by a significant suppression of the associated neuroinflammation, as indicated by decreased IBA1 immunoreactivity in both the cortex and hippocampus (Fig. 6f,g and Extended Data Fig. 7l,m).

We sought to determine if this widespread mitigation of converging pathologies translated into tangible neuroprotection. We performed NeuN immunostaining to quantify surviving neurons in the hippocampus, a region severely affected in this model. Ir nanozyme-treated mice showed significantly higher NeuN-positive cell counts in both the hippocampal cornu ammonis 1 (CA1) and dentate gyrus (DG) subregions compared to the vehicle control group, indicating that the Ir nanozyme effectively preserved neuronal integrity amidst a complex pathological assault (Fig. 6h,i). Collectively, these results demonstrate that the Ir nanozyme can disrupt the cycle of cross-pathology seeding between Aβ, tau, and αSyn. This highlights its potential as a multi-target therapeutic strategy for complex neurodegenerative disorders characterized by the convergence of multiple proteinopathies, such as AD with Lewy bodies.

## Conclusions

In this study, by developing a pair of nanozymes with opposing RONS-modulating capabilities, we created a unique *in vivo* condition to ascertain the importance of RNS in αSyn pathology. The Ir□Cu nanozyme which promotes NO•, dramatically exacerbated αSyn propagation despite being a superior ROS scavenger. Mechanistically, we show that this Ir□Cu-driven RNS promotion triggers the downstream DNA damage-PARP-1 activation cascade, a pathway previously linked to αSyn neurodegeneration^6^. The ability of a NOS inhibitor, L-NAME, to completely abrogate this entire cascade—from PAR accumulation to the final pS129 pathology—provides evidence that excess RNS, independent of bulk ROS levels, is sufficient to drive αSyn propagation. This work does not necessarily arbitrate a universal “ROS vs. RNS” debate, but rather provides causal evidence of the pathogenic potential of the RNS driving the PAR pathway in a complex *in vivo* setting.

Our findings on copper’s potent ability to modulate RNS-driven synucleinopathy may offer a new lens through which to view the impact of environmental risk factors on neurodegeneration^56^. Excessive copper exposure can result from ingestion of dissolved copper in drinking water^57,58^, use of copper-containing fungicides and fertilizers^58^, or inhalation of copper-laden atmospheric particulate matter^58,59^. Growing evidence has suggested a link between environmental copper exposure and neurodegenerative diseases^31,32,60^. For instance, a case report documented early-onset Lewy-body dementia in association with chronic exposure to contaminated drinking water by copper leaching from household plumbing^30^. In addition, we and others have recently highlighted the strong association between fine particulate matter (PM_2.5_) air pollution—a significant component of which is primarily derived from human activities—and the risk for LBD and PD^61,62^. Although ambient copper concentrations in outdoor air are generally low^59^, atmospheric copper—mainly originating from traffic-related sources such as brake and tire wear—has been identified as a major contributor to the oxidative potential of atmospheric particulate matter^63^. Notably, elevated levels of airborne copper, along with other metal species, have been detected in specific indoor environments, such as subway systems^64–66^. However, the health effects of metal-rich PM_2.5_ in these settings remain poorly understood^67,68^.

Furthermore, our study provides a critical insight for the broader field of nanomedicine. The paradoxical, pathology-exacerbating effect of the Ir□Cu nanozyme serves as a cautionary tale: the design of therapeutic nanozymes, especially those containing redox-active metals like copper^33^, must be approached with caution. The potential for unintended catalytic activities could inadvertently fuel the very pathological processes they are designed to treat, underscoring the absolute necessity of comprehensive mechanistic and safety profiling in nanozyme development.

Finally, and in contrast to the risks posed by selective RNS promotion, our work validates a powerful therapeutic strategy based on neutralizing RONS. The copper-free Ir nanozyme, by effectively scavenging both ROS and the pathogenic RNS pathway, demonstrated substantial therapeutic efficacy. Its success across diverse and challenging preclinical models—including brain-first, body-first/gut-first, and AD/PD co-pathology paradigms—not only provides strong validation for this broad-spectrum RONS-scavenging pathway but also establishes this class of nanozyme as a highly promising therapeutic candidate for a broad spectrum of α-synucleinopathies.

In summary, our work leverages a rationally designed nanozyme probe to uncover the importance of RNS-driven PAR-related pathogenesis in α-synucleinopathy. This discovery highlights the critical role of nitrosative stress in driving αSyn propagation; it offers a plausible mechanistic basis for the neurotoxicity linked to environmental copper exposure; and it serves as a crucial design principle for nanomedicine, mandating caution in the use of redox-active metals for neuroprotective applications. Ultimately, by demonstrating the efficacy of a broad-spectrum RONS-scavenging nanozyme, our study validates this neutralization approach as a therapeutic strategy for Parkinson’s disease and related α-synucleinopathies.

## Methods

### Chemicals

Iridium(III) chloride (IrCl_3_), L-Ascorbic acid (AA) and 3,3′,5,5′-tetramethylbenzidine dihydrochloride (TMB), Copper(□) chloride dihydrate (CuCl_2_•2H_2_O), terephthalic acid (TA), polyvinylpyrrolidone (PVP, MW = 30000), sodium hydroxide (NaOH), hydrogen peroxide (H_2_O_2_), Iron(□) sulfate heptahydrate (FeSO_4_•7H_2_O), and hydrochloric acid (HCl) were purchased from were purchased from Sinopharm Chemical Reagent Co., Ltd (Shanghai, China). S-nitroso-N-acetylpenicillamine (SNAP) was obtained from the Shanghai Aladdin Biochemical Technology Co., Ltd. (Shanghai, China). 2-(4-Carboxyphenyl)-4,5-dihydro-4,4,5,5-tetramethyl-1H-imidazol-1-yloxy-3-oxide potassium salt (PTIO) was obtained from the Shanghai Maokang Biotechnology Co., Ltd (Shanghai, China). Diethylenetriaminepentaacetic acid (DTPA), xanthine oxidase (XOD) and xanthine (HX) were purchased from Sigma-Aldrich (Shanghai, China). 5,5-Dimethyl-1-pyrroline N-oxide (DMPO) and 5tert-Butoxycarbonyl-5-methyl-1-pyrroline-N-oxide (BMPO) were purchased from DOJINDO (Shanghai, China). Chemicals sources are listed in Supplementary Table 4. Milli-Q water (18 MΩ cm) was used in the preparation of all solutions and all reagents were not further purified before using.

### Synthesis of Ir and Ir-Cu nanoalloys

Taking the preparation of Ir_3_Cu nanozyme as an example, 440 mg polyvinyl pyrrolidone (PVP, MW = 30000), 0.1127 g of L-Ascorbic acid (AA), 3 mL of 20 mM IrCl_3_, and 1 mL of 20 mM CuCl_2_ were added to a beaker filled with 6.6 mL of H_2_O, stirred and sonicated for 5 minutes to uniformly disperse them. The solution was transferred to a 25 mL Teflon-lined stainless-steel autoclave and heated at 200 °C for 6 hours before it was cooled to room temperature. The product was precipitated with acetone (1:3 by volume) to remove impurities, and then deionized water was added to the product to a constant volume of 8 mL for subsequent use. Other nanozymes preparation methods with different proportions are basically the same, as long as the molar ratio of Ir^3+^/Cu^2+^ is changed, and the total concentration of metal ions in the solution remains unchanged. Pure Ir nanozymes are prepared by the same procedure except for without addition of Cu^2+^.

### Characterization

The crystal structures of the Ir and Ir-Cu alloy nanoparticles were characterized by X-ray diffraction (XRD, D8 Advance diffractometer, Bruker, Germany) using monochromatized Cu Kα radiation (λ = 1.5418 Å). Transmission electron microscopy (TEM) images were captured on a Tecnai G2 F20 U-TWIN electron microscope (FEI, USA) with an accelerating voltage of 200 kV and an FEI Titan HRTEM microscope (FEI, USA) with an accelerating voltage of 300 kV. Elemental mapping and line scan was verified by scanning transmission electron microscope (STEM) and energy dispersive X-ray (EDS) spectroscopy in the high angle annular dark field (HAADF) mode. Infrared spectra were obtained using a Nicolet 6700 Fourier Transform Infrared (FTIR) Spectrometer (Thermo Fisher Scientific, USA). UV-vis-NIR absorption spectra were obtained using a Cary 5000 UV-vis-NIR Spectrometer (Varian, USA) and a matched quartz cuvette with a path length of 1 cm. Electron spin resonance (ESR) spectra were obtained using an EMX micro electron paramagnetic resonance spectrometer (Bruker, Germany).

### Peroxidase-like activities assays

The TMB assay was used to testing peroxidase-like activity^69^. Typically, 20 μL of 20 mM TMB solution, 20 μL of 0.1M H_2_O_2_ and 3 mL water were mixed in a 1 cm thick quartz cuvette, then different types of Ir-Cu nanozymes was added to initiate the oxidation of TMB. The UV-visible absorption spectra were recorded immediately at selected time intervals for the calculation of the oxidation rate of TMB. All experiments in this test are repeated at least 3 times. Unless otherwise noted, reactions were performed at room temperature. The POD-like activity was detected by the same method for other various mono metal NPs, CDs and Cu based nanozymes (bimetallic alloy and CDs-Cu).

### Catalase-like activities assays

For testing catalase-like activity, a dissolved oxygen electrode is used. Add 300 μL of 100 mM H_2_O_2_ to a small beaker containing 5 mL PBS buffer (10 mM, pH 7.4) and mix well as a working solution. Add different types of Ir-Cu nanozymes to the working solution to trigger the reaction, stir slowly with magnetons, and monitor the dissolved oxygen content in the working solution with a dissolved oxygen electrode, reading the value at 0.5 min intervals. All experiments in this test are repeated at least 3 times. The CAT-like activity was detected by the same method for other various mono metal NPs, CDs and Cu based nanozymes (bimetallic alloy and CDs-Cu).

### Test for the ability to scavenge superoxide

To test the O_2_^•-^ free radical scavenging, using two methods.

1. ESR direct detection of O ^•-70^:15 μL H_2_O, 10 μL 5 mM hypoxanthine, 10 μL 0.25 mM TDPA, 5 μL 250 mM BMPO and different types of Ir-Cu nanozymes were added to the centrifuge tube and mixed evenly. Add another 5 μL of 0.4 U/mL XOD and mix well (total volume 50 μL). Seal with a glass capillary with a 50 μL inner diameter of 1 mm and insert it into the ESR cavity to record the spectra at the selected time. The SOD-like activity was detected by the same method for other various mono metal NPs, CDs and Cu based nanozymes (bimetallic alloy and CDs-Cu). The condition for ESR measurements in detection of the spin adducts BMPO/•OOH: 20 dB microwave power attenuation, 1 G field modulation, 100 G scanning range, and 2 mW microwave power for spin adduct detection.
2. UV-vis absorption spectrometry: 0.6 mL 1.75 mM 18-crown-6 (DMSO solution) and different types of Ir-Cu nanozymes were added to the dry and clean cuvette, then 70 μL 25 mM KO_2_ (DMSO solution) was added to the dry and clean cuvette, quickly mixed well, waited for 3 min. Then, add 2.4 mL of 0.33 mM NBT (aqueous solution) and mix well, wait for 2 min, and measure UV-vis. All experiments in this test are repeated at least 3 times.

### Test for the ability to scavenge **•**OH

To test the •OH free radical scavenging, using two methods.

1. ESR direct detection •OH^71^: 20 μL 10 mM PBS, 10 μL 100 mM H_2_O_2_, 5 μL DMPO and different types of Ir-Cu nanozymes were added to the centrifuge tube and mixed evenly. Then add 10 μL 10 mM Fe^2+^ and mix well (total volume 50 μL). Seal with a glass capillary with an inner diameter of 50 μL inner diameter of 1 mm and insert it into the ESR cavity, and record the spectra after 5 min of reaction. The •OH scavenging activity was detected by the same method for other various mono metal NPs, CDs and Cu based nanozymes (bimetallic alloy and CDs-Cu). The condition for ESR measurements: 10 mW microwave power; 100 G scan range and 1 G field modulation.
2. Fluorescence spectroscopy: add 2 mL of 0.5 mM terephthalic acid (TA) and 50 μL of 100 mM H_2_O_2_ to the centrifuge tube containing 1 mL H_2_O and mix well, then different types of Ir-Cu nanozymes and 50 μL 10 mM Fe^2+^ to mix well, and wait for 4 hours. After 4 h, perform a fluorescence test scan with an excitation wave of 315 nm. The fluorescence spectroscopy test conditions for other different nanozymes are basically the same.

### Test for the ability to scavenge or promote NO•

The NO• scavenging activity of nanozymes were tested by ESR. NO• was produced by S-nitroso-N-acetylpenicillamine (SNAP) and was trapped by PTIO to form typical ESR signals. For the ESR measurement, 5 µL SNAP (2mM) was mixed with 5 µL PTIO (2 µM) in pH = 7.4 PBS buffer in the absence and presence of different types of Ir-Cu nanozymes. Seal with a glass capillary tube with an inner diameter of 1 mm and a volume of 50 µL, insert it into the ESR chamber, and record the spectrum at the selected reaction time intervals. The condition for ESR measurement: 10 mW microwave power; 100 G scan range and 1G field modulation. The ability to scavenge or promote NO• was detected by the same method for other various mono metal NPs, CDs and Cu based nanozymes (bimetallic alloy and CDs-Cu).

### Fluorescent labeling of nanozymes

Nanozyme were incubated with Cy5.5 PEG Thiol (3.4k Da, Nanocs, New York, NY, USA) in Milli-Q water at 4 °C overnight. After labeling, the mixture was washed three times with Milli-Q water and purified by centrifugation at 14,000□rpm for 40□min each time.

### Animals

5XFAD Tg mice (Tg6799 line) overexpress known familial AD (FAD) mutations in APP (K670N/M671L [isoform770] + I716V + V717I) and PS1 (M146L + L286V) genes, under the control of the Thy1 promoter. 5XFAD mice were bred by crossing male Tg mice with female B6/SJL F1 mice. Original 5XFAD and all B6/SJL breeders were purchased from the Jackson Laboratory (Bar Harbor, ME; 5XFAD: Stock No: 034840-JAX; B6/SJL: Stock No: 100012). A53T mice (Tg(Prnp-SNCA*A53T)23Mkle, JAX Stock No:004479) expressing human α-synuclein with the A53T mutation under the mouse prion promoter and wild-type C57BL/6J mice (Stock No: 000664,) were obtained from the Jackson Laboratory. Mice were housed in a temperature-controlled room under a 12-hour light/dark cycle with free access to food and water. All procedures were performed in accordance with the NIH Guide for the Care and Use of Experimental Animals. Studies were approved by the Institutional Animal Care and Use Committee of John Hopkins University.

### Cell cultures

Primary cortical neurons were isolated from embryonic day 15.5 (E15.5) CD1 mouse (Charles River Laboratories, Wilmington, MA, USA) embryos as previously described^8^. Pregnant dams were euthanized, and embryos were rapidly removed under sterile conditions. The cerebral cortices were dissected in ice-cold Hank’s Balanced Salt Solution (HBSS, without calcium and magnesium; Thermo Fisher Scientific, Waltham, MA, USA), and the meninges were carefully removed. Tissues were then enzymatically dissociated in 0.25% trypsin-EDTA (Thermo Fisher Scientific) at 37 □ for 15 minutes, followed by gentle mechanical trituration. The resulting single-cell suspension was passed through a 70 μm cell strainer, centrifuged at 320g for 5 minutes, and resuspended in Neurobasal medium supplemented with 10% B-27 (Thermo Fisher Scientific), 0.5 mM L-glutamine, and 1% penicillin-streptomycin. Cells were seeded onto poly-D-lysine (PDL, 50 μg/mL)-coated plates at a density of 1 × 10□ cells/cm² and maintained at 37□°C in a humidified incubator with 5% CO□. Half of the culture medium was replaced every 3–4 days. Experiments were typically performed on day 7 in vitro.

Primary microglia were isolated from the cerebral cortices of postnatal day 1 (P1) CD1 mouse pups (Charles River Laboratories) as previously described^8,72^. Brains were rapidly dissected under sterile conditions, and the meninges were carefully removed. Cortical tissue was minced and digested in 0.25% trypsin-EDTA (Thermo Fisher Scientific) at 37 °C for 15 minutes. Following enzymatic digestion, the tissue was gently triturated using Pasteur pipettes and passed through a 100□μm cell strainer to obtain a single-cell suspension. Cells were seeded into poly-D-lysine (PDL, 50□μg/mL)-coated T75 flasks and cultured in DMEM/F12 medium (Gibco) supplemented with 2 mM L-glutamine, 100 μM non-essential amino acids, 2 mM sodium pyruvate 10% fetal bovine serum (FBS, Thermo Fisher Scientific) and 1% penicillin-streptomycin. Cultures were maintained at 37□°C in a humidified incubator with 5% CO□, and the medium was replaced every 5 days. After 10–14 days in vitro, microglia were isolated by magnetic cell sorting using the EasySep™ Mouse CD11b Positive Selection Kit (StemCell Technologies, Vancouver, BC, Canada), following the manufacturer’s protocol. The purity of isolated microglia was confirmed by IBA1 immunostaining.

HEK-A53T-YFP cells^49,50^, a human embryonic kidney 293 cell line (HEK293) engineered to express a familial Parkinson’s disease-associated αSyn mutant (A53T) fused to yellow fluorescent protein (YFP) were cultured in DMEM supplemented with 10% FBS and 1% penicillin-streptomycin, and maintained at 37□°C in a humidified 5% CO_2_ incubator.

### **α**Syn PFF preparation

Recombinant mouse αSyn protein was expressed and purified as previously decribed^73^. PFF were generated by incubating αSyn monomers in PBS buffer (5 mg/mL) with constant agitation (1,000 rpm, 37 °C) for 7 days, followed by sonication for 1 minute (1□second on, 1□second off) at 30% amplitude using a Branson Digital Sonifier (Branson Ultrasonics, Danbury, CT, USA).

### RONS assay

Intracellular ROS detection: Mouse primary microglia were treated with αSyn PFF (20 μg/mL) and nanozymes (Ir, Ir_3_Cu, IrCu, or IrCu_3_; 12.5 μM) for 12 hours. After treatment, cells were incubated with CM-H□DCF-DA (5□μM; C6827; Thermo Fisher Scientific) at 37□°C for 30□minutes to assess intracellular ROS levels. Fluorescence signals were analyzed by co-staining with Hoechst (1:5000, 62249, Thermo Fisher Scientific) following image by confocal microscopy. ImageJ software was used for fluorescence quantification.

Intracellular NO• detection: primary microglia were treated with αSyn PFF (20 μg/mL) and nanozymes (Ir, Ir_3_Cu, IrCu, or IrCu_3_; 12.5 μM) for 12 hours. Intracellular NO• levels were measured using DAF-FM diacetate (5□μM; D23844, Thermo Fisher Scientific) at 37°C for 30□minutes. Fluorescence signals were detected by confocal microscopy and quantified using ImageJ.

### Cellular internalization and biodistribution

HEK-A53T-YFP cells were incubated with Cy5.5-labeled Ir nanozymes (12.5□μM) for 12 hours. Following incubation, cells were washed with PBS, fixed with 4% paraformaldehyde (PFA) in PBS, and imaged using fluorescence microscopy to visualize the intracellular localization of the Ir nanozymes. Fluorescence intensity was quantified using ImageJ software.

*In vivo* biodistribution and brain enrichment: Cy5.5-labeled Ir nanozymes (5□mM in 100□μL PBS) were administered to 3-month-old male C57BL/6 wild-type (WT) mice via either intranasal (*i.n.*) or intravenous (*i.v.*) injection. Three days post-administration, hair over the skull was carefully removed, and whole-body fluorescence imaging was performed using the IVIS Spectrum Imaging System (PerkinElmer, Waltham, MA, USA) to assess brain accumulation. Mice were subsequently euthanized, and major organs—including the brain, heart, liver, spleen, lung, kidney, and whole blood—were collected for *ex vivo* fluorescence imaging to evaluate the biodistribution of the Ir nanozymes.

Gastrointestinal Tract Enrichment: To assess gastrointestinal retention of Ir nanozymes, 3-month-old male A53T mice were orally gavaged with Cy5.5-labeled Ir nanozymes (5□mM in 200□μL PBS). Three days post-gavage, the gastrointestinal tract was isolated, and luminal contents were flushed out with PBS. Fluorescence imaging was then performed using the IVIS Spectrum Imaging System (PerkinElmer).

### PFF injection

Stereotaxic striatal PFF injection: Three-month-old male C57BL/6J mice were anesthetized with a ketamine-xylazine cocktail and positioned in a stereotaxic apparatus. Mice received a unilateral injection of αSyn PFF (5□μg in 2□μL PBS) into the dorsal striatum using a 5□μL Hamilton syringe equipped with a 30-gauge needle. The stereotaxic coordinates relative to bregma were: anteroposterior (AP) +0.2 mm, mediolateral (ML) +2.0 mm, dorsoventral (DV) −2.6 mm.

Stereotaxic hippocampal PFF injection: Nine-month-old female 5XFAD mice were deeply anesthetized with a ketamine-xylazine cocktail and secured in a stereotaxic frame. αSyn PFF (5□μg in 2□μL PBS) or PBS (control) was injected unilaterally at a rate of 0.2□μL/minute using a Hamilton syringe. The injection was targeted to the left dorsal hippocampus (bregma: −2.5 mm; ML: +2.0 mm; DV: −2.4 mm), followed by the overlying cortex (DV: −1.4 mm), as the needle was gradually withdrawn.

Intestinal intramuscular PFF injection: Three-month-old male A53T mice were anesthetized and maintained on a heating pad to preserve core body temperature. The abdominal cavity was opened to expose the stomach, and a total of 25□μg of α-synuclein preformed fibrils (αSyn PFF) in 5□μL PBS was injected into the wall of the pyloric stomach and intestine wall of the duodenum (0.5 cm apart from the pyloric stomach injection site) using a 10□μL Hamilton syringe as previously described^54^. Injections were made near the myenteric plexus. Following injection, the abdominal incision was sutured, and mice were returned to standard housing conditions for post-operative recovery.

### Nanozyme treatments *in vivo*

To test in vivo effects of nanozymes in the brain-first PD model, starting two days after stereotaxic striatal injection of αSyn PFF, mice received weekly treatments for two months with vehicle or nanozymes (Ir or Ir□Cu; 10□mM in 100□μL), administered either *i.n.* or *i.v.*. At two months post-PFF injection, mice were euthanized, and brains were harvested for immunofluorescence analysis.

For behavioral evaluation, a separate cohort of mice received weekly intranasal administration of Ir nanozyme (10□mM in 100□μL) for 4 months, beginning two days after stereotaxic striatal injection of αSyn PFF. Behavioral tests were performed at nine months post-PFF injection. Following behavioral assessments, mice were euthanized, and brains and spleens were collected for immunohistochemical analysis.

In a separate experimental group, beginning two days after striatal PFF injection, mice received daily intraperitoneal injections of the nitric oxide synthase inhibitor L-NAME (80□mg/kg; N5751, Millipore Sigma), in combination with daily intranasal administration of Ir□Cu nanozyme (10□mM in 20□μL) for one month. One-month post-injection, mice were euthanized, and brain tissues were collected for immunofluorescence analysis and RNA isolation for subsequent RNA sequencing.

To investigate the therapeutic effect of Ir nanozyme in a peripheral-origin PD model, Ir nanozyme (10□mM in 200□μL) was administered by oral gavage weekly for two months, starting two days after Intestinal intramuscular PFF injection. At the endpoint, brain and gastrointestinal tissues were collected for immunofluorescence analysis.

To evaluate the therapeutic efficacy of Ir nanozyme in a model of Alzheimer’s disease with coexisting Lewy body pathology, Ir nanozyme (10□mM in 100□μL) or vehicle was administered intranasally once per week for three months, starting two days after stereotaxic hippocampal PFF injection. Brains were collected at three months post-injection for pathological analysis.

### Immunofluorescence and immunohistochemistry analysis

Immunofluorescence staining of primary neurons: Primary neurons were fixed with 4% PFA in PBS for 15 minutes at room temperature, followed by permeabilization with 0.2% Triton X-100 in PBS for 10 minutes. After washing, neurons were incubated overnight at 4°C with primary antibodies diluted in blocking buffer (10% normal goat serum in PBS), including: anti-NeuN (1:50, MAB377, Sigma-Aldrich, St Louis, MO, USA) and anti-phospho-αSyn (pS129; 1:1000, ab51253, Abcam, Cambridge, MA, USA). The following day, cells were washed and incubated with appropriate fluorophore-conjugated secondary antibodies for 1 hour at room temperature. Antibody sources and dilutions are listed in Supplementary Table 5. Nuclei were counterstained with Hoechst, and images were acquired using a confocal microscope and analyzed using ImageJ.

Immunofluorescence staining of tissue sections: coronal brain sections or intestinal sections were permeabilized with 0.2% Triton X-100, and blocked with 10% normal goat serum for 30 minutes. Sections were then incubated overnight at 4□°C with primary antibodies, including: anti-IBA1 (1:1000, 019-19741, Fujifilm Wako), anti-3-NT (1:50, ab61392; Abcam), anti-4-HNE, (1:200, MA527570, Thermo Fisher Scientific); anti-PAR (1:500)^6^; anti-γH2A.x (1:500, 05-636, Millipore Sigma); anti-AT8 (1:500, MN1020, Thermo Fisher Scientific), anti-Aβ_42_ (1:500, 010-26903, Fujifilm Wako), and anti-NeuN (1:50, MAB377, Millipore Sigma). For intestinal myenteric plexus staining, sections were stained with antibodies against IBA1, pS129, and tyrosine hydroxylase (TH; 1:500, AB152, Sigma-Aldrich, St. Louis, MO, USA). The following day, sections were washed and incubated with appropriate fluorophore-conjugated secondary antibodies for 1 hour at room temperature. Nuclei were counterstained with Hoechst. Images were acquired using a confocal microscope and analyzed using ImageJ.

Immunohistochemistry: Free-floating sections were blocked using Avidin/Biotin Blocking System (927301, BioLegend, San Diego, CA, USA) and 10% goat serum in PBS with 0.2% Triton X-100. Sections were then incubated with rabbit anti-TH antibody (1:1000, AB152), followed by biotinylated goat anti-rabbit IgG secondary antibody. Signal was amplified using streptavidin-conjugated horseradish peroxidase (HRP) from the VECTASTAIN® ABC Kit (Vector Laboratories, Burlingame, CA, USA) and visualized using SigmaFast DAB peroxidase substrate (Sigma-Aldrich). Sections were counterstained with Nissl dye (FD Thionin Solution™ Double Strength, PS101-02, FD NeuroTechnologies, Columbia, MD, USA). The number of TH-positive and Nissl-positive neurons in the substantia nigra pars compacta were quantified using unbiased stereological methods and their projections in the striatum were quantified by ImageJ.

### Behavioral tests

To evaluate the therapeutic effects of Ir nanozymes on αSyn PFF-induced behavioral deficits, mice injected with PBS or αSyn PFF and treated with Ir nanozymes were subjected to a battery of behavioral tests. Motor function was assessed using the pole test, rotarod test, and grip strength test. Additionally, open field test was conducted to evaluate general locomotor activity and anxiety-like behavior.

#### Pole test

To assess motor coordination and bradykinesia, mice were habituated in the behavior testing room for 30 minutes prior to testing. The apparatus consisted of a vertical metal pole (75□cm in height, 9□mm in diameter) wrapped in surgical gauze to aid grip. Each mouse was placed head-up near the top of the pole (7.5□cm from the peak), and the total time taken to descend to the base was recorded. Animals underwent training for two consecutive days, with three trials per session. On the test day, the maximum trial duration was limited to 60 seconds. Parameters recorded included time to turn and total time.

#### Rotarod

To evaluate motor coordination and balance, mice were pre-trained on a rotarod for three consecutive days. On the fourth day, animals were placed on a rotating rod that accelerated from 4 to 40 rpm over 5 minutes. The latency to fall (or to passively cling without walking for two full turns) was measured. Each mouse underwent three trials, and the average latency time was used for analysis.

#### Grip strength test

Forelimb neuromuscular strength was assessed using a grip strength meter (BIO-GS3, Bioseb, FL, USA). Each mouse was allowed to grasp a metal grid with its forelimbs. Gentle traction was applied by pulling the tail horizontally until the animal released its grip. The peak force exerted before release was recorded digitally and expressed in grams (g). Three measurements per animal were taken, and the average was used for analysis.

#### Open field

Spontaneous locomotion and anxiety-like behavior were assessed in a square open-field chamber (40 × 40 × 40□cm), subdivided into 36 equal squares (6 × 6). The central area was defined as the four innermost squares, and the remaining area constituted the periphery. Mice were placed in the center and allowed to freely explore for 5 minutes under dim light conditions. The arena was cleaned with 10% ethanol between animals. A video tracking system (ANY-maze software) was used to quantify time spent in the central zone-parameters indicative of anxiety-related behavior.

### Human RNA-Seq data processing

We obtained raw FASTQ files from a previous study that performed bulk RNA sequencing of anterior cingulate cortex samples from 13 PD patients and 6 non-neurological control samples. We processed the FASTQ files using the SPEAQeasy^74,75^ RNA-seq pipeline, which integrates several bioinformatics tools to convert raw FASTQ files into ready-to-use R objects for differential gene expression analysis. In brief, reads were initially quality-checked with FastQC^72^ v0.11.8, followed by trimming with Trimmomatic^76^ v0.39 when necessary. The quality-checked reads were then aligned to the human reference genome (hg38/GRCh38) using HISAT2^77^ v2.2.1. Finally, gene-level expression was quantified using featureCounts^78^ based on GENCODE release 25 annotations.

### Mouse RNA sequencing analysis

RNA was extracted from the cortex tissues encompassing the anterior cingulate cortex (ACC) region of mice treated with Veh, Ir_3_Cu nanozyme, or Ir_3_Cu nanozyme with L-NAME, with three replicates per condition. RNA isolation was performed using TRIzol reagent (Thermo Fisher Scientific) followed by purification with the RNeasy Mini Kit (QIAGEN), according to the manufacturers’ instructions. Directional mRNA library was prepared using poly (A) enrichment by Novogene Corporation Inc. (Sacramento, CA, USA). The libraries were constructed using a strand-specific protocol and sequenced on the Illumina NovaSeq X Plus platform, generating 150 bp paired-end reads (PE150) with a minimum sequencing depth of 12 Gb per sample. The RNA-seq data were processed using the same SPEAQeasy RNA-seq pipeline, with reads aligned to the mouse genome (mm10) and gene-level expression quantified based on GENCODE release M25. Lowly expressed genes were first filtered out by retaining genes with at least 10 reads in at least 3 samples. The filtered data was then normalized using the trimmed mean of M-values (TMM) method, implemented in the calcNormFactors function from the edgeR^79^ package. Principal component analysis (PCA) was performed on the mouse RNA-seq data using the top 1000 most variable genes, determined by variation of expression levels measured as log_2_ normalized counts per million (CPM) across samples.

To assess sample quality and detect potential outliers, we applied a network-based approach using Z-connectivity statistics. Specifically, after TMM normalization, log_2_-transformed CPM values were used to compute pairwise biweight midcorrelations (bicor) between samples. A signed adjacency matrix was then constructed using the adjacency() function from the WGCNA package^80^, representing the overall similarity between samples. For each sample, connectivity was calculated and converted into a standardized Z-score. One sample from the PFF group had a Z-score close to the defined threshold (Z = -3) and was flagged as an outlier; it was excluded from the downstream differential expression analysis.

### Differential gene expression analysis

Differential gene expression analysis for both human and mouse RNA-seq datasets was conducted using the DESeq2^81^ package. For the human RNA-seq data, we adjusted for potential confounders in the regression model, including sex, age at death, post-mortem interval (PMI), and RNA integrity number (RIN). We only included condition status for the mouse data in the regression model to detect group difference of gene expression. To enhance visualization and ranking of genes, we applied the apeglm method^82^ for effect size (Log_2_FC) shrinkage implemented in DESeq2.

### Gene set enrichment analysis

Gene Set Enrichment Analysis (GSEA) was conducted using the clusterProfiler^83^ package, focusing on gene sets from Gene Ontology (GO) related to biological processes. We set the minimum gene set size (minGSSize) to 20 and the maximum gene set size (maxGSSize) to 500. The input for GSEA was a ranked gene list derived from the shrunken effect sizes obtained with DESeq2.

### Comparison of transcriptional responses in mice and human patients

We compared the transcriptional responses in mice under each condition to those in human patients with PD based on a set of 14,066 homologous genes common to both species. Following a previously published approach^84^, comparison was limited to genes that showed differential expression in humans, defined by a *P*-value < 0.05 and an absolute log_2_(fold change) greater than log_2_(1.2). To ensure a sufficient number of genes for cross-species comparison, we used raw (unshrunken) effect sizes from DESeq2. The strength of correlation was assessed using Spearman correlation.

#### Quantification and statistical analysis

All data (except epidemiological and transcriptomic data) were analyzed using GraphPad Prism 8. Statistics Data are presented as the mean ± SEM (Standard Error of the Mean) or presented as violin plots showing all individual data points. With at least 3 independent experiments. Representative morphological images were obtained from at least 3 experiments with similar results. Statistical significance was assessed via a *t*-test, one way ANOVA test, or two-way ANOVA test followed by indicated post-hoc multiple comparison analysis. Assessments with *P* < 0.05 were considered significant.

## Acknowledgements

The authors acknowledge the joint participation by the Adrienne Helis Malvin Medical Research Foundation through its direct engagement in the continuous active conduct of medical research in conjunction with The Johns Hopkins Hospital and the Johns Hopkins University School of Medicine and the Foundation’s Parkinson’s Disease Program M-2023. T.M.D. is the Leonard and Madlyn Abramson Professor in Neurodegenerative Diseases.

## Author contributions

X.M, W.H, T.M.D conceived the idea, led the project, designed experiments and supervised the discussion, wrote and edited the manuscript. S.H. processed RNA-seq data analysis, wrote and edited the manuscript. J.H., V.L.D. and P.L. designed experiments and supervised the discussion, wrote and edited the manuscript. S.L., Y.M., N.W., J.Z., X.Z., Y.C., J.S., Q.Y., R.K., D.J., J.Y.S., X.Y., A.K., J.P., Y.G., S.Y., S.Z. conducted all the experiments, data curation and analysis, and interpretation of nanozyme, wrote the original draft. B.G.K. provided PAR antibody, R.C. provided PFF.

---

**Extended Data Fig. 1.**
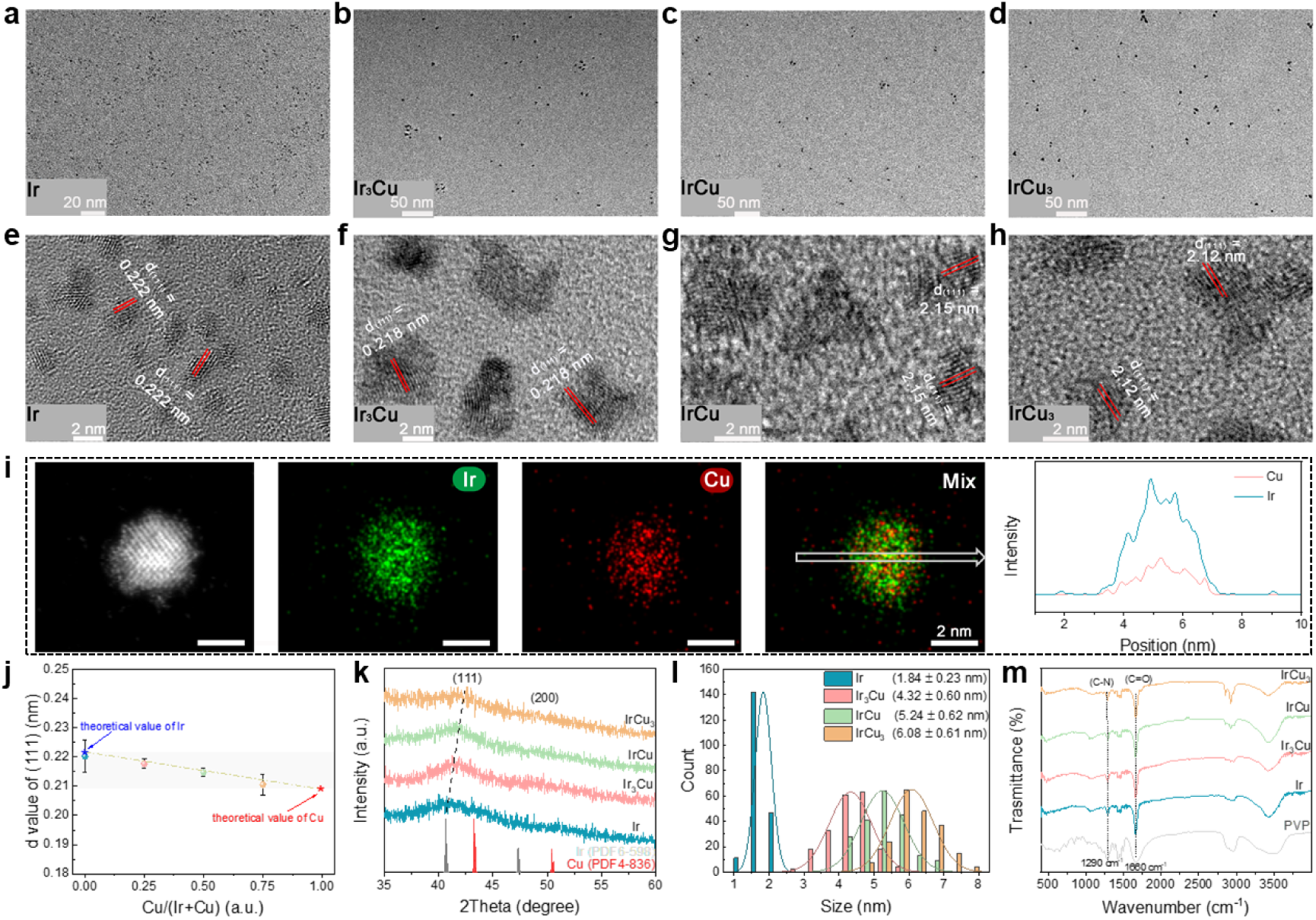
Structure characterization for the Ir and Ir-Cu nanozymes with different Ir/Cu ratio. **a**-**d**, TEM images of Ir (**a**), Ir_3_Cu (**b**), IrCu (**c**) and IrCu_3_ (**d**), respectively. **e**-**g**, HRTEM images of Ir (**e**), Ir_3_Cu (**f**), IrCu (**g**) and IrCu_3_ (**h**), respectively. The spacing between the two red parallel lines represents the lattice spacing. **i**, HAADF-STEM image and corresponding EDS mapping and line scan of Ir and Cu elements for Ir_3_Cu. The white box-shaped arrows represent the position of the line scan. **j**, The linear fitting of the lattice spacing of Ir and Ir-Cu nanozymes with different Ir/Cu ratios to the Cu/(Ir+Cu) ratio. The lattice spacing measured from the HRTEM image for the Ir and Ir-Cu nanozymes with different Ir/Cu ratio. Data are presented as mean ± SEM, *n* = 3. **k**, XRD patterns of Ir, Ir_3_Cu, IrCu, and IrCu_3_, along with the standard PDF card of Ir and Cu, inserted for comparison. **l**, Size distribution of Ir, Ir_3_Cu, IrCu and IrCu_3_. The data measured from the TEM images were randomly counted with 200 values for each sample. **m**, FT-IR spectra of Ir, Ir_3_Cu, IrCu, IrCu_3_ and PVP.

**Extended Data Fig. 2.**
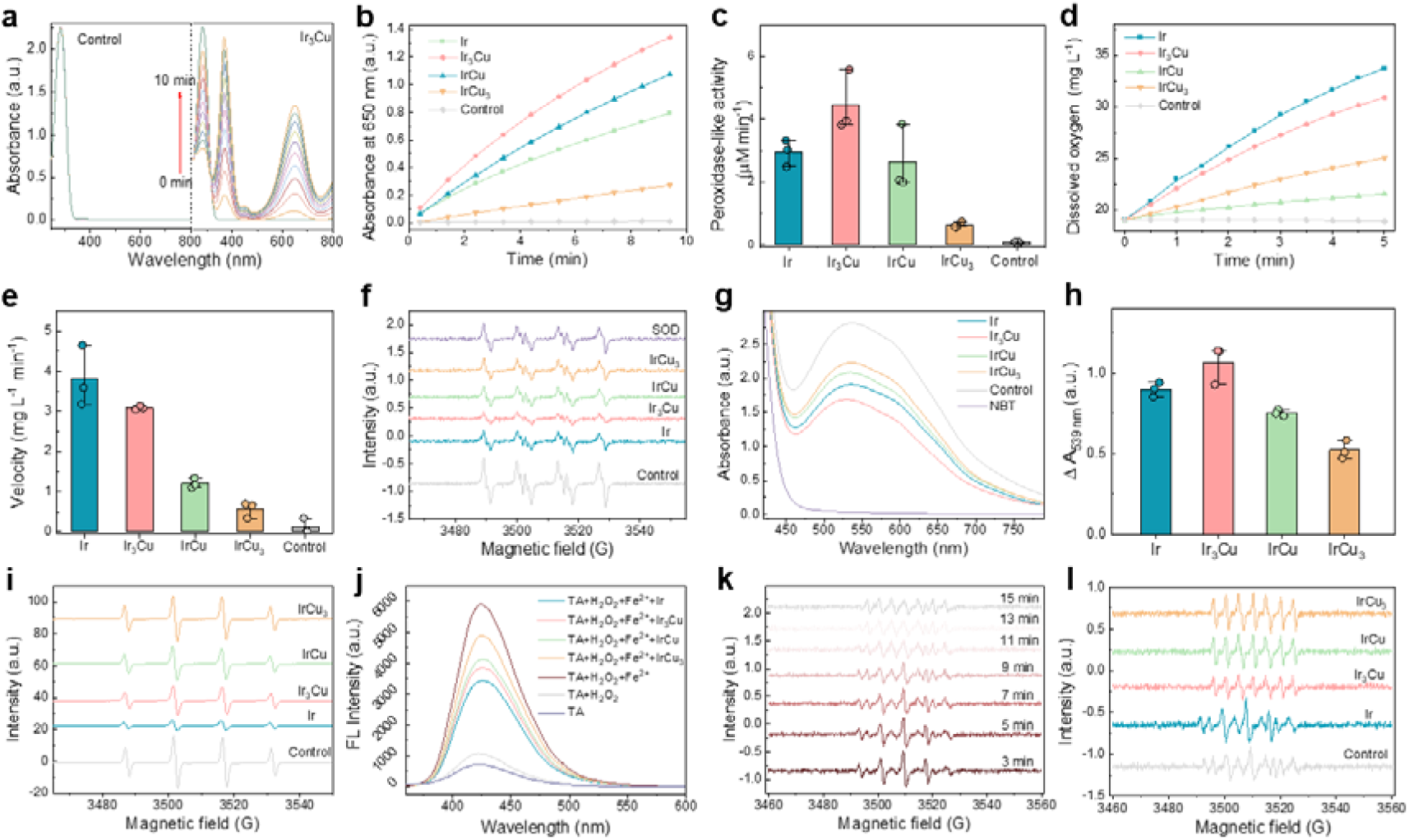
The ROS and NO• regulating capacity of multi-enzyme like Ir and Ir-Cu nanozymes with different Ir/Cu ratio. **a**, UV-vis spectra evolution with time for solutions containing TMB and H_2_O_2_ in the presence or absence of IrCu_3_. **b**, The absorbance changes at 650 nm as a function of reaction time during TMB oxidation catalyzed by Ir and Ir-Cu nanozymes with different Ir/Cu ratio in the presence of H_2_O_2_. **c,** A comparison of the POD-like activity of Ir and Ir-Cu nanozymes with different Ir/Cu ratio. Data are presented as mean ± SEM, *n* = 3. **d**, Evolution of oxygen content in solution with time decomposed H_2_O_2_ by Ir and Ir-Cu nanozymes with different Ir/Cu ratio. **e**, A comparison of the CAT-like activity of Ir and Ir-Cu nanozymes with different Ir/Cu ratio. Data are presented as mean ± SEM, *n* = 3. **f**, ESR spectra of BMPO/O_2_^•-^ generated from the samples containing XOD, Xanthine, BMPO, and different samples Ir and Ir-Cu nanozymes with different Ir/Cu ratio. **g**, UV-vis spectra of NBT method for scavenging O_2_^•-^ by Ir and Ir-Cu nanozymes with different Ir/Cu ratio. **h,** A comparison of the SOD-like activity of Ir and Ir-Cu nanozymes with different Ir/Cu ratio by NBT method. Data are presented as mean ± SEM, *n* = 3. **i**, ESR spectra of DMPO/•OH generated from the samples containing DMPO, H_2_O_2_, Fe^2+^, and with different samples of Ir and Ir-Cu nanozymes with different Ir/Cu ratio. **j**, FL spectra of TA solution by Ir and Ir-Cu nanozymes with different Ir/Cu ratio for •OH scavenging. **k**, Evolution of ESR spectra of PTIO/NO• generated from the samples containing PTIO and SNAP over time. **l**, ESR spectra of PTIO/NO• generated from the samples containing PTIO, SNAP, and in the absence (control) and presence of Ir and Ir-Cu nanozymes with different Ir/Cu ratio. Extended Data Figure 2 compares systematically the Ir and Ir-Cu alloy nanozymes with different Ir/Cu ratios for their ability to regulate ROS/RNS. Kinetic evaluation of POD-like activity via TMB oxidation assays revealed Ir_3_Cu nanozymes exhibit the highest initial reaction rate (IRR) (Extended Data Fig. 2a,b), with activity trends following: Ir > Ir_3_Cu > IrCu > IrCu_3_ (Extended Data Fig. 2c). Further mechanistic studies demonstrated time-, H_2_O_2_ concentration-, and pH-dependent catalytic behavior for Ir_3_Cu nanozymes. Michaelis-Menten analysis using TMB and H_2_O_2_ as substrates yielded apparent *K*_m_ (57.14 μM) and *V*_max_ (1407.77 μM·s^-1^), confirming enhanced substrate affinity relative to natural horseradish peroxidase. In addition, alloy composition profoundly influenced CAT-like activity, with pure Ir nanozymes demonstrating maximal oxygen evolution rates and IRR (Extended Data Fig. 2d,e), followed by Ir_3_Cu > IrCu > IrCu_3_ in descending order. SOD-mimetic capacity, quantified through ESR spectroscopy and NBT reduction assays (Extended Data Fig. 2f–h), showed all nanozymes effectively scavenged O_2_^−•^ radicals, with efficiency ranking: Ir_3_Cu > Ir > IrCu > IrCu_3_. The O_2_^−•^ scavenging efficiency was promoted with the addition of Ir_3_Cu nanozymes rising from 5 μL to 15 μL, which proved the SOD-like activity to be concentration-dependent. Both ESR and fluorescein-based assays further revealed •OH radical scavenging capabilities in Fenton reaction systems (Fe^2+^/H_2_O_2_) (Extended Data Fig. 2i,j), with activity trends: Ir > Ir_3_Cu > IrCu > IrCu_3_. Nitric oxide free radical (NO•) is a representative RNS and recognized as a detrimental cause for PDs. To test the interaction of Ir and Ir-Cu nanozymes with NO•, SNAP was selected as a NO• generator and PTIO was chosen as a spin label molecule. PTIO is a typical NO• scavenger that reacts with NO• to produce ESR-silent PTI, leading to gradual reduction of the five-line spectrum and formation of a seven-line signal (Extended Data Fig. 2k). Interestingly, Ir nanozymes efficiently scavenged NO• a dosage of 3 μg/mL eliminated the NO• signal while Ir-Cu nanozymes with different compositions accelerated NO• production (Extended Data Fig. 2l).

**Extended Data Fig. 3.**
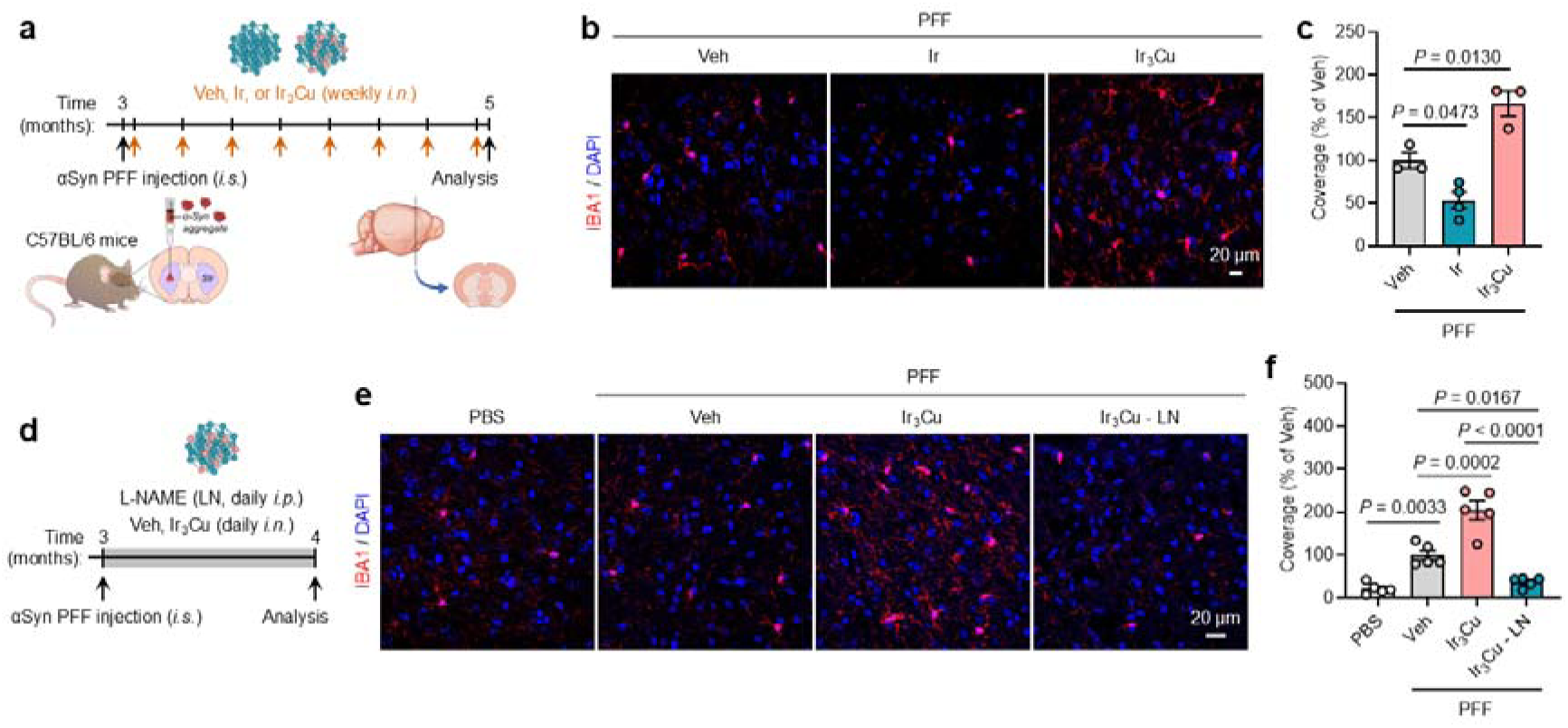
Intranasal nanozyme treatment modulates microglial activation in αSyn PFF-injected mice. **a**, Schematic showing C57BL/6 wide-type mice received intrastriatal injection (*i.s.*) of αSyn PFF, followed by weekly *i.n.* Veh or nanozymes (Ir or Ir_3_Cu). Brains were collected 2 months after PFF injection. **b**, Representative images and (**c**) corresponding quantification of IBA1 positive microglia in the cortex, with coverage normalized to Veh group. *n* = 3–4 mice per group. **d**, Schematic showing C57BL/6 mice received *i.s.* αSyn PFF injection, followed by *i.n.* Veh or Ir_3_Cu along with intraperitoneal (*i.p.*) L-NAME injection. Brains were collected 1-month post-injection. **e**, Representative images and (**f**) corresponding quantification of IBA1 positive microglia in the cortex, with coverage normalized to Veh group. *n* = 5 mice per group. Data are presented as mean ± SEM. *P* < 0.05 by one-way ANOVA with Tukey’s multiple comparisons test.

**Extended Data Fig. 4.**
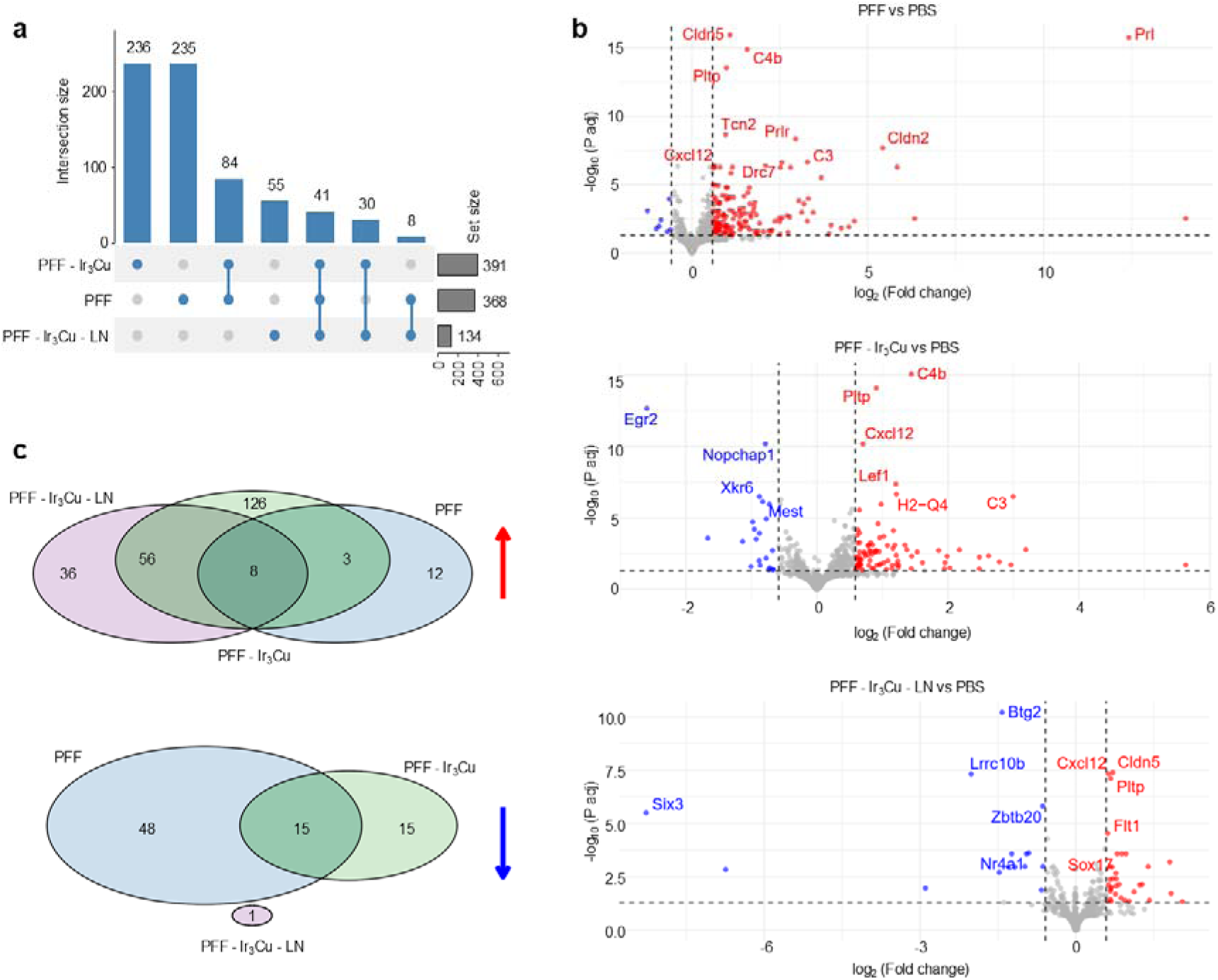
Transcriptomic responses to PFF, PFF-lr_3_Cu, and PFF-lr_3_Cu-LN in WT mice. **a**, UpSet plot illustrating the number of differentially expressed genes (DEGs) unique to or shared across the three conditions. **b**, Volcano plots for each condition, highlighting top DEGs based on statistical significance and effect size. **c**, Venn diagrams of significantly enriched gene sets for each treatment. Upregulated gene sets (top, red arrow) are predominantly immune-related, while downregulated gene sets (bottom, blue arrow) are mostly involved in neuronal functions.

**Extended Data Fig. 5.**
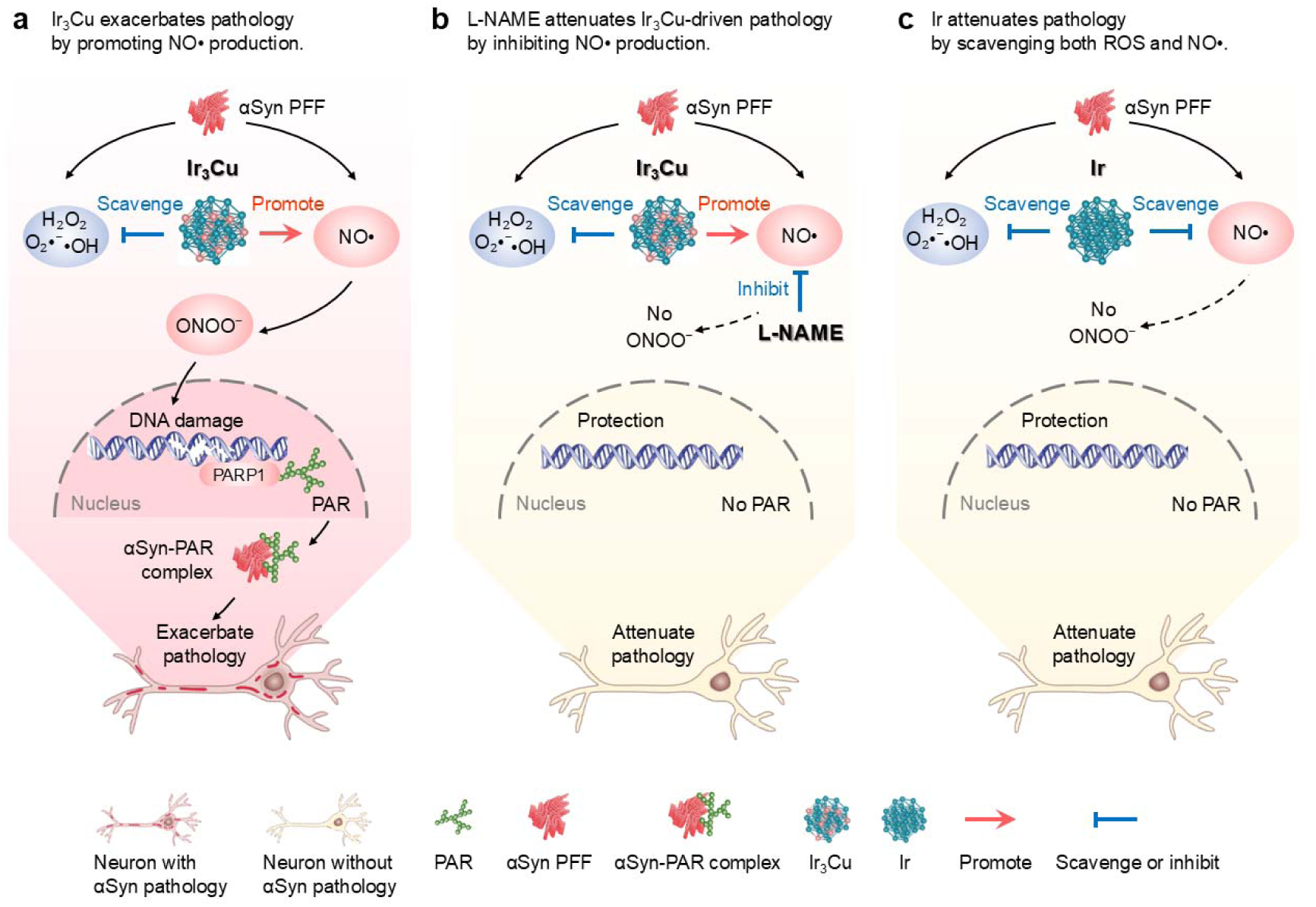
Divergent modulation of RONS by Ir and Ir_3_Cu nanozymes in α-Syn pathology and its impact on disease progression. **a–c,** In αSyn PFF-induced PD mice: **a**, Ir_3_Cu nanozyme selectively scavenges reactive oxygen species (ROS; e.g., H_2_O_2_, O_2_^•^□, •OH) while promoting NO• production, thereby facilitating peroxynitrite (ONOO□) formation. This cascade triggers DNA damage, PARP1 activation, PAR production, and the formation of PAR-αSyn aggregates complexes, ultimately exacerbating PD pathology; **b**, L-NAME inhibits Ir_3_Cu nanozyme-induced NO• production and prevents subsequent neurotoxic events; **c**, In contrast, Ir nanozyme scavenges both ROS and NO•, suppressing ONOO□ formation, attenuating DNA damage, and protecting against NO•-mediated αSyn pathology.

**Extended Data Fig. 6.**
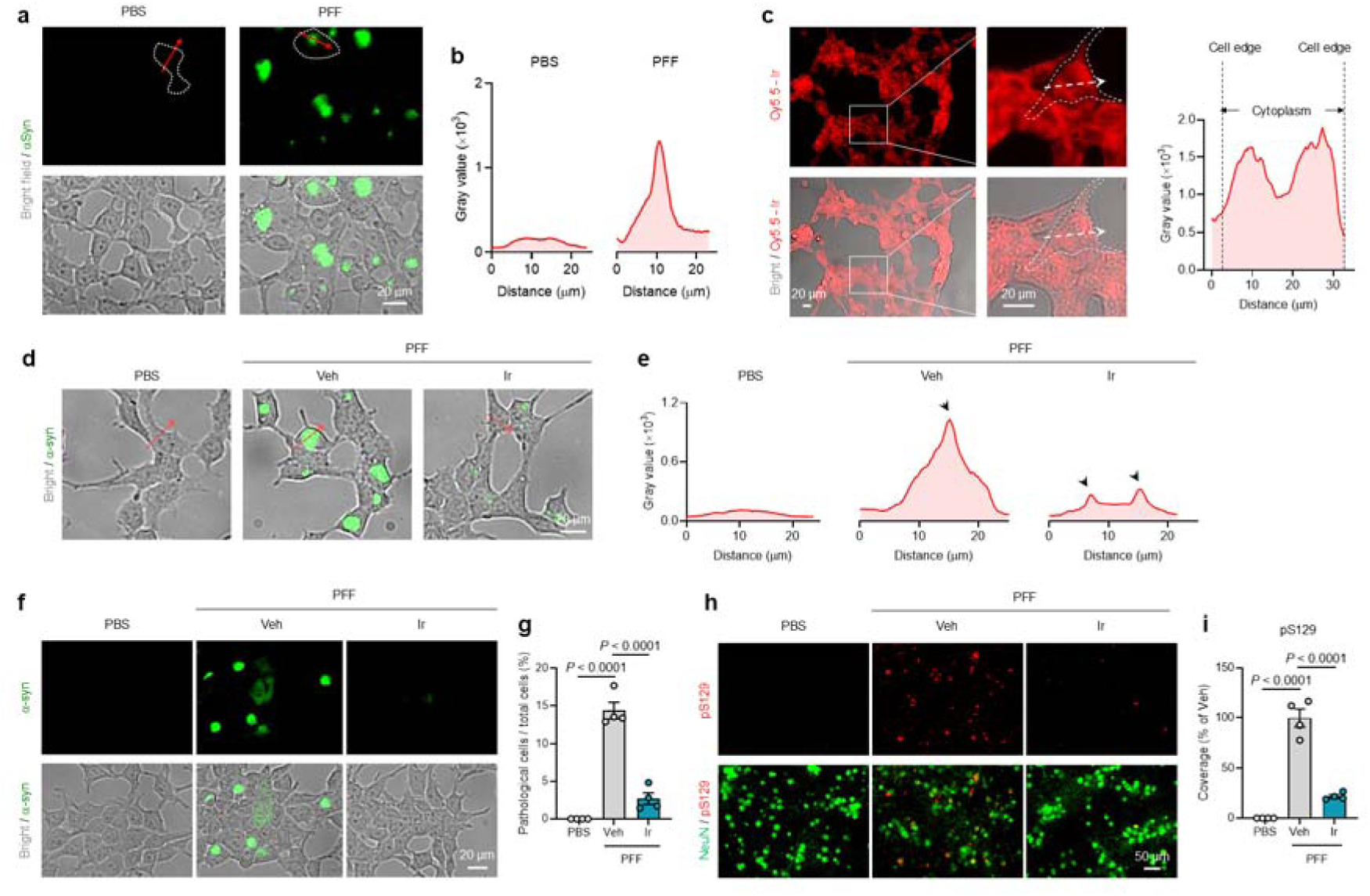
Ir nanozyme inhibits αSyn aggregates and pathology *in vitro*. **a**, Representative images displaying HEK-A53T-YFP cells with αSyn aggregates after treated with αSyn PFF. **b**, The plot profiles of YFP fluorescence, with red line-arrows indicating the regions analyzed. **c**, Representative images showing the cellular internalization of Cy5.5-labelled Ir in HEK-A53T-YFP cells. The plot profiles of Cy5.5 fluorescence intensity along the red arrows indicated in (**c**). **d**,**e**, HEK-A53T-YFP cells were treated with αSyn PFF (5□μg/mL) at 3 hours prior to Ir nanozyme administration (125□μM). After 48 hours, cells were imaged to detect intracellular αSyn aggregates. **d**, Representative images and (**e**) corresponding plot profiles of YFP fluorescence, with red arrows indicating the regions analyzed. **f**,**g**, HEK-A53T-YFP cells were co-treated with αSyn PFF (5□μg/mL) and Ir nanozyme (125□μM) for 48 hours. **f**, Representative images showing αSyn aggregation in HEK-A53T-YFP cells. **g**, Quantification of the percentage of cells displaying αSyn aggregates (pathologic cells) relative to the total number of cells. *n* = 4. **h**, Representative images and (**i**) corresponding quantification of pS129 (red) in NeuN (greed)-positive mouse primary neurons treated with αSyn PFF and Ir nanozyme, with coverage normalized to the Veh group. *n* = 4. Data are presented as mean ± SEM, *P* < 0.05 by one-way ANOVA with Tukey’s multiple comparisons test.

**Extended Data Fig. 7.**
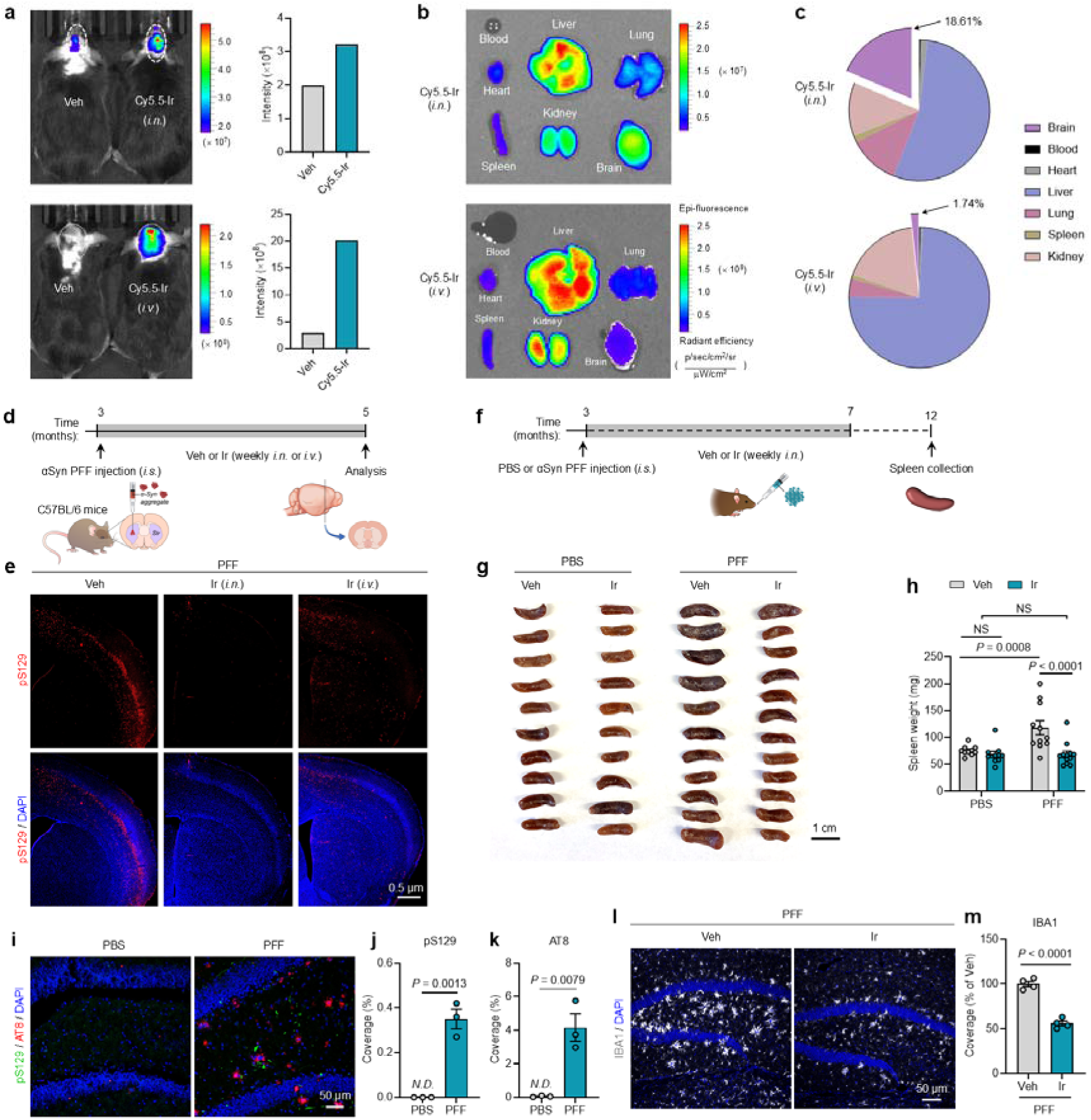
Biodistribution of Ir nanozyme and evaluation of administration routes for therapeutic efficacy. **a,** IVIS Spectrum imaging showing brain enrichment distribution of Cy5.5 labelled Ir (Cy5.5-Ir) following either intranasal (*i.n.*) or intravenous (*i.v.*) administration, and corresponding quantification of Cy5.5-Ir fluorescence intensity in the brain. **b**, *Ex vivo* IVIS images showing the biodistribution of Cy5.5-Ir in the major organs—including the brain, blood, heart, liver, lung, spleen, and kidney—following administration *via i.n.* and *i.v* injection. **c**, Quantification of biodistribution of Cy5.5-Ir in the brain, blood, heart, liver, lung, spleen, and kidney following *i.n* or *i.v.* administration. **d**, Schematic showing C57BL/6 wide-type mice received intrastriatal injection of αSyn PFF, followed by weekly *i.n.* or *i.v.* administration of Veh or Ir nanozyme. Brains were collected 2 months post-PFF injection. **e**, Immunofluorescence images of pS129-positive αSyn pathology (red) in brain sections of PFF-injected mice treated with Ir nanozyme *via* either *i.n.* or *i.v.* route. **f**, Schematic showing C57BL/6 WT mice received *i.s.* αSyn PFF or PBS injection, followed by weekly *i.n.* administration of Ir nanozyme for 4 months. Spleens were collected 9 months post-PFF injection. **g**, Representative images of spleens collected from each treatment group, along with (**h**) quantification of spleen weights to assess peripheral immune activation. *n* = 10–12 mice per group Data are presented as mean ± SEM. *P* < 0.05 by two-way ANOVA with Tukey’s multiple comparisons test. **i**, Representative images of pS129 (green) and AT8 p-tau (red) in the hippocampus of 5XFAD mice following intra-hippocampal injection of PBS or αSyn PFF. **j**,**k**, Corresponding quantification of (**j**) pS129-positive αSyn pathology and (**k**) AT8-positive tau pathology, with coverage normalized to the Veh group. *n* = 3 mice per group. **l**, Representative images of IBA1-positive microglia in the hippocampus and (**m**) corresponding quantification of the coverage normalized to Veh group. *n* = 4 mice per group. Data are presented as mean ± SEM. *P* < 0.05 by unpaired two-tailed Student’s *t*-test.

