## Supplementary material for "Copper-containing ROS-scavenging nanozyme paradoxically drives α-synucleinopathy by amplifying nitrosative stress"

**Table of contents**

**Supplementary Fig. 1 |** Gene set enrichment analysis of PFF-Ir_3_Cu *vs* PBS control.

**Supplementary Fig. 2 |** Gene set enrichment analysis of PFF-Ir_3_Cu-LN *vs* PBS control.

**Supplementary Fig. 3 |** Gene set enrichment analysis of PFF *vs* PBS control.

**Supplementary Table 1 |** Gene set enrichment analysis of PFF-Ir_3_Cu *vs* PBS control.

**Supplementary Table 2 |** Gene set enrichment analysis of PFF-Ir_3_Cu-LN *vs* PBS control.

**Supplementary Table 3 |** Gene set enrichment analysis of PFF *vs* PBS control.

**Supplementary Table 4 |** The list of chemicals used in this study.

**Supplementary Table 5 |** The list of antibodies used in this study.

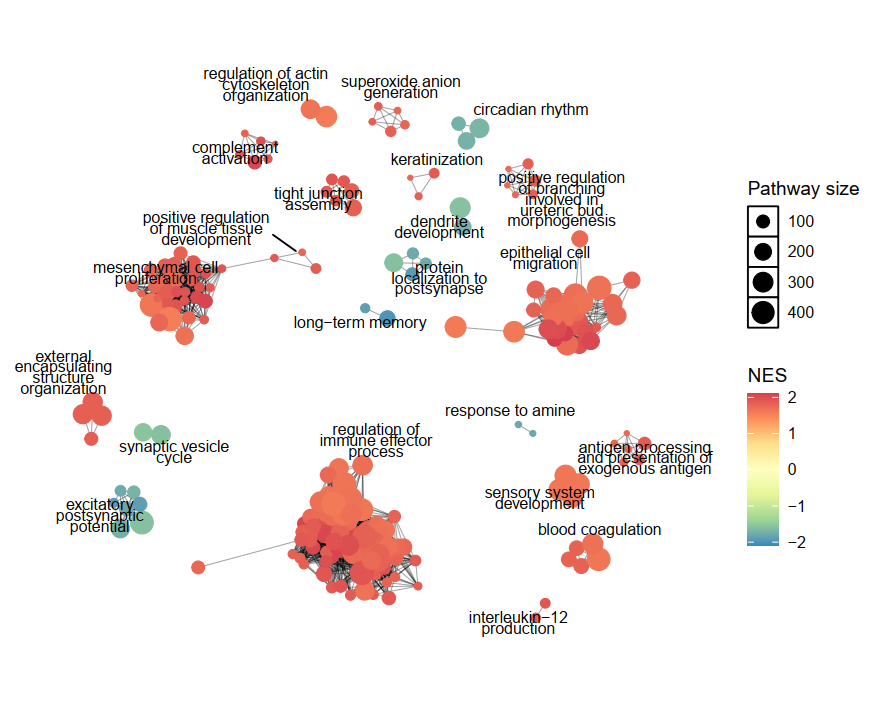

**Supplementary Fig. 1 |** Gene set enrichment analysis of PFF-Ir_3_Cu *vs* PBS control.

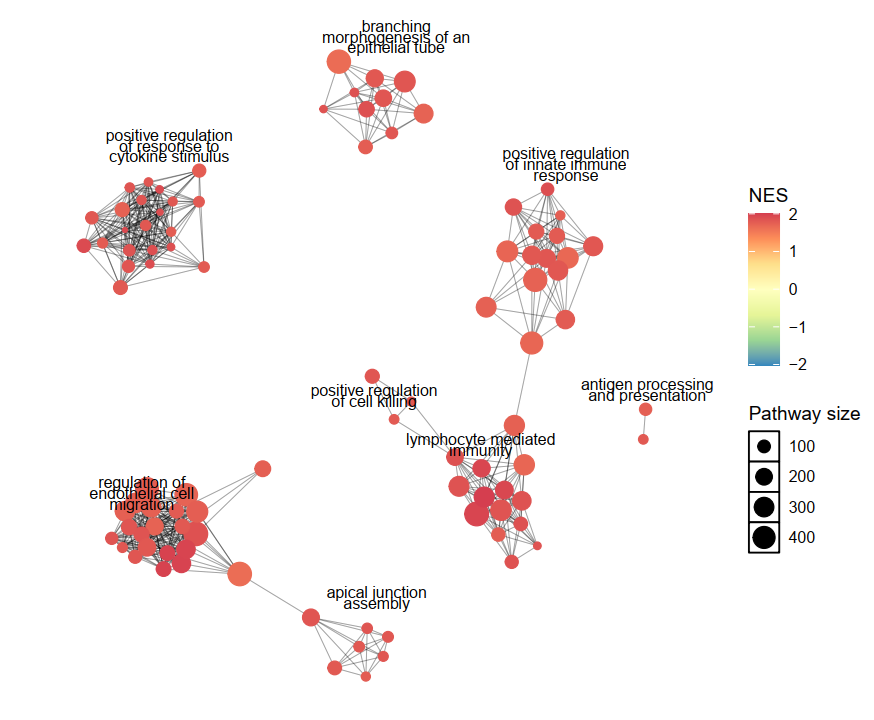

**Supplementary Fig. 2 |** Gene set enrichment analysis of PFF-Ir_3_Cu-LN *vs* PBS control.

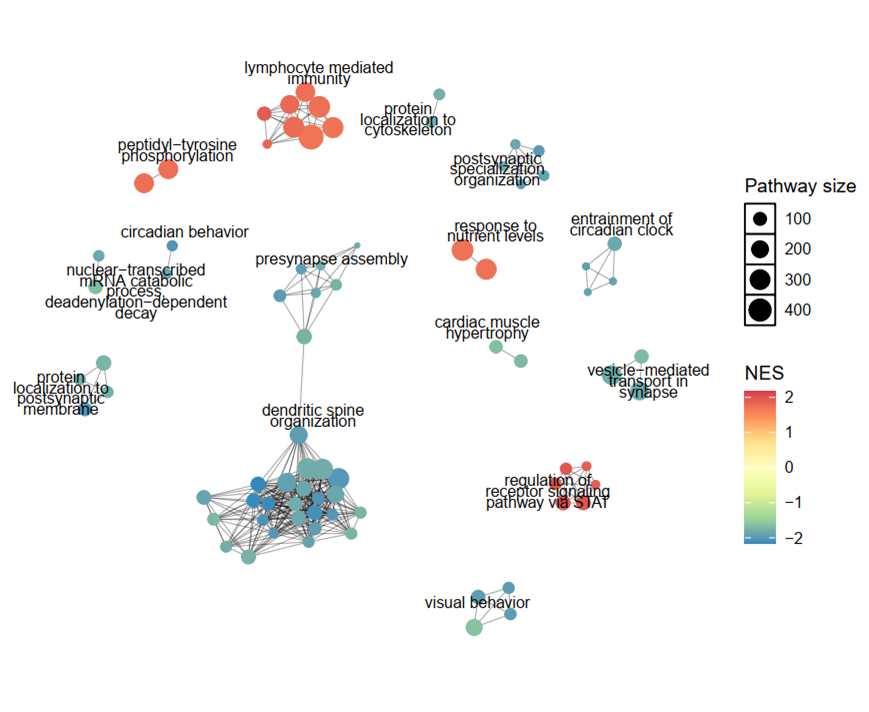

**Supplementary Fig. 3 |** Gene set enrichment analysis of PFF *vs* PBS control.

**Supplementary Table 1 |** Gene set enrichment analysis of PFF-Ir_3_Cu *vs* PBS control.

**Supplementary Table 2 |** Gene set enrichment analysis of PFF-Ir_3_Cu-LN *vs* PBS control.

**Supplementary Table 3 |** Gene set enrichment analysis of PFF *vs* PBS control.

**Supplementary Table 4 |** The list of chemicals used in this study.

| **REAGENT or RESOURCE** | **SOURCE** | **IDENTIFIER** |
| --- | --- | --- |
| **Chemicals** | | |
| Iridium(III) chloride （**IrCl_3_**，≥99.80%） | Sinopharm Chemical Reagent | CAS：10025-83-9 |
| L-Ascorbic acid （**AA**，≥99.70%） | Sinopharm Chemical Reagent | CAS：50-81-7 |
| 3,3′,5,5′-tetramethylbenzidine dihydrochloride （**TMB**, ≥99.00%） | Sinopharm Chemical Reagent | CAS：64285-73-0 |
| Copper(Ⅱ) chloride dihydrate (**CuCl_2_·2H_2_O**, ≥99.00%) | Sinopharm Chemical Reagent | CAS：10125-13-0 |
| Terephthalic acid (**TA**, ≥99.00%) | Sinopharm Chemical Reagent | CAS：100-21-0 |
| Polyvinylpyrrolidone K30 (**PVP**, ≥99.80%) | Sinopharm Chemical Reagent | CAS：9003-39-8 |
| Sodium hydroxide (**NaOH**, ≥99.70%) | Sinopharm Chemical Reagent | CAS：1310-73-2 |
| Hydrogen peroxide (**H_2_O_2_**, ≥30.0%) | Sinopharm Chemical Reagent | CAS：7722-84-1 |
| Iron(Ⅱ) sulfate heptahydrate (**FeSO_4_·7H_2_O**, 99.0～101.0%) | Sinopharm Chemical Reagent | CAS：7782-63-0 |
| Hydrochloric acid (**HCl**, 36.00～38.00%) | Sinopharm Chemical Reagent | CAS：7647-01-0 |
| (S)-Nitroso-N-acetylpenicillamine (**SNAP**, ≥97.00%) | Shanghai Aladdin Biochemical | CAS：67776-06-1 |
| 2-(4-Carboxyphenyl)-4,5-dihydro-4,4,5,5-tetramethyl-1H-imidazol-1-yloxy-3-oxide potassium salt (**PTIO**, ≥98.00%) | Shanghai Maokang Biotechnology | CAS：148819-94-7 |
| Diethylenetriaminepentaacetic acid (**DPTA**, ≥98.00%) | Sigma-Aldrich | CAS：67-43-6 |
| Xanthine oxidase (**XOD**, ≥0.4 units/mg protein ) | Sigma-Aldrich | CAS：9002-17-9 |
| Xanthine (**C_5_H_4_N_4_O_2_**, ≥99.00%) | Sigma-Aldrich | CAS：69-89-6 |
| 5,5-Dimethyl-1-pyrroline N-oxide (**DMPO**, ≥99.00%) | DOJINDO | CAS：3317-61-1 |
| 5-tert-Butoxycarbonyl-5-methyl-1-pyrroline-N-oxide (**BMPO**, ≥99.00%) | DOJINDO | CAS：B568-10 |

**Supplementary Table 5 |** The list of antibodies used in this study.

| **REAGENT or RESOURCE** | **SOURCE** | **IDENTIFIER** | **Dilution** |
| --- | --- | --- | --- |
| **Antibodies** | | |  |
| α-Synuclein (phosphor S129) (pS129; clone: EP1536Y) | Abcam | ab51253 | 1:1000 |
| NeuN (clone: A60) | Sigma-Aldrich | MAB377 | 1:50 |
| Ionized calcium-binding adapter molecule1 (IBA1; clone: BC05) | Fujifilm Wako | 019-19741 | 1:1000 |
| 3-nitrotyrosine (3-NT; clone: 39B6) | Abcam | ab61392 | 1:50 |
| 4-hydroxynonenal (4-HNE; clone: 12F7) | Thermo Fisher Scientific | MA5-27570 | 1:200 |
| Poly(ADP-ribose) (PAR) | Made in Dawson lab | N/A | 1:500 |
| Phospho-Histone H2A.X (γH2A.x; clone: JBW301) | Millipore Sigma | 05-636 | 1:500 |
| Phospho-Tau (Ser202, Thr205) (AT8; clone: AT8) | Thermo Fisher Scientific | MN1020 | 1:500 |
| Amyloid β_42(43)_ (Aβ_42(43)_; clone: BC05) | Fujifilm Wako | 010-26903 | 1:500 |
| Tyrosine Hydroxylase (TH) | Sigma-Aldrich | AB152 | 1:500 |
